# Identification of Region-Specific Gene Isoforms in the Human Brain Using Long-Read Transcriptome Sequencing and Their Correlation with DNA Methylation

**DOI:** 10.1101/2023.05.13.540603

**Authors:** Mihoko Shimada, Yosuke Omae, Akiyoshi Kakita, Ramil Gabdulkhaev, Taku Miyagawa, Makoto Honda, Akihiro Fujimoto, Katsushi Tokunaga

## Abstract

**Background:** Site specificity is known in neuropsychiatric disorders, and differences in gene expression patterns could potentially explain this mechanism. However, studies using long-read transcriptome sequencing to analyze gene expression in different regions of the human brain have been limited, and none have focused on the hypothalamus, which plays a crucial role in regulating autonomic functions.

**Results:** We performed long-read RNA sequencing on 12 samples derived from three different brain regions of the same individuals; the cerebellum, hypothalamus, and temporal cortex. We found that, compared to other regions, many genes with higher expression levels in the cerebellum and temporal cortex were associated with neuronal pathways, whereas those with higher expression levels in the hypothalamus were primarily linked to immune pathways. In addition, we investigated genes with different major isoforms in each brain region, even with similar overall expression levels among regions, and identified several genes, such as *GAS7*, that express different major isoforms in different regions. Many of these genes are involved in “actin filament-based process” and “cell projection organization” pathways, suggesting that region-dependent isoforms may have distinct roles in dendritic spine and neuronal formation in each region. Furthermore, we investigated the involvement of DNA methylation in these isoforms and found that DNA methylation may be associated with isoforms that have different first exons.

**Conclusions:** Our results provide potentially valuable findings for future research on brain disorders and shed light on the mechanisms underlying isoform diversity in the human brain.

## Background

Brain disorders such as neurodegenerative diseases and psychiatric disorders are multifactorial diseases with complex etiologies that contribute significantly to human morbidity and mortality. Site specificity is known in neuropsychiatric disorders, and different neurodegenerative diseases and their clinical symptoms are associated with distinctive neurodegenerative systems [1]. For example, in amyotrophic lateral sclerosis, the upper and lower motor neurons in the motor cortex, brain stem nuclei and anterior horn of the spinal cord are impaired [2]. In frontotemporal lobar degeneration, the pregenual anterior cingulate cortex (ACC) to the midcingulate cortex are affected, whereas Alzheimer disease involves posterior cingulate regions and the ACC is unaffected [1]. Although pathology studies suggested the involvement of several factors in this site specificity, such as changes in neurotransmitters, protein homeostasis and energy demand, the detailed mechanism is still unclear, and other changes that are not measurable by histological techniques, such as synaptic connections or gene expression patterns, may also be involved [1]. Previous studies on psychiatric disorders examined gene expression in postmortem brain samples of patients with schizophrenia and bipolar disorder. These studies identified altered expression of GABA-related genes in the superior temporal gyrus and hippocampus in both disorders [3–5] and anterior cingulate cortex-specific expression changes in schizophrenia [6]. Identifying and understanding the specific genes and gene expression patterns in each brain regions may provide crucial insights into the underlying mechanisms of various neurological and psychiatric diseases.

Previous studies on gene expression analysis in the brain have revealed not only genes involved in brain disorders, but also transcripts that are altered by factors such as sleep status and aging [7–9]. However, most of these analyses were based on short-read sequence or microarray data, and their isoforms were only inferred from sequences with short read-lengths. Recent advances in transcriptome technology using long-read sequencing allow us to read the entire length of the transcript from the 5’ to the 3’ UTR and polyA tail at once, enabling us to capture more complex and precise transcript features without phasing problems [10]. Previous studies have reported that tissue-specific protein-coding and non-coding transcripts are more abundant in the brain compared to other organs [11], and that their expression is also elevated overall [12]. The brain transcriptome is considered to be relatively specific, and it is expected that new findings will be obtained by using full-length transcriptome analysis to accurately evaluate individual isoforms involved in the alternative splicing in the brain. A recent study analyzed the single-cell long reads of cells from various regions of the postmortem mouse brain, and reported region-specific expression of isoforms, which is very important in unraveling the complex integration of cells in the brain [13]. While studies using animal models and cells are invaluable, many studies have investigated human brain site-specific isoforms [7, 12, 14–18], and evaluating postmortem human brain samples is considered essential in investigations of the complex biology of the human brain and causes of brain diseases. Indeed, numerous novel isoforms and human-specific gene expression patterns have been detected in studies that have analyzed postmortem human brain samples using long-read methods [19, 20]. However, these studies focused on the differences in the expression of the cerebral cortex between mice and humans, changes in the expression of the cerebral cortex during developmental stages, and provided an overview of gene expression in various tissues including the brain. As a result, the findings of these studies do not provide a lot of information on the regional specificity of the brain.

Here, we performed long-read sequencing of RNA extracted from three regions of the postmortem human brain –– the temporal cortex, hypothalamus, and cerebellum –– with a focus on regional specificity. The temporal cortex is involved in language, memory, and hearing [21–23], and has been reported to be associated with schizophrenia [24], Alzheimer disease [25] and temporal lobe epilepsy [26]. The hypothalamus is a critical brain region that regulates endocrine and autonomic functions [27, 28]. It is composed of numerous neuronal nuclei and coordinates and manages diverse physiological functions, including thermoregulation, the stress response, feeding behavior, and sleep arousal. The hypothalamus has been also a target for the treatment of heterogeneous psychopathological symptoms [27, 29]. The cerebellum is primarily involved in the integration of sensory and motor functions and is associated with several developmental disorders [30, 31]. The purpose of this study was to elucidate the region-dependent gene and isoform expression patterns in the human brain using full-length transcript sequencing and to obtain data that can be used as a control for future disease research. We analyzed three regions of the postmortem brains of four male subjects who had no brain lesions or infections and died in their 50s or later. We attempted to minimize the influence of genomic sequences and other environmental factors by making comparisons among different regions from the same samples. In addition, epigenetics may play a significant role in the intra-individual expression differences. It is well known that the methylation of CpG islands in the promoter region typically results in silencing of gene expression, and that gene-body methylation is associated with higher levels of gene expression [32–34]. DNA methylation has been also reported to be involved in alternative splicing by acting as an indicator of an alternative intra-genic promoter [35], or by recruiting methyl CpG binding protein 2 (MeCP2) to promote exon recognition [36, 37]. Despite such evidence, the relationship between DNA methylation and alternative splicing is not fully understood, and very few studies have evaluated the contribution of DNA methylation to alternative splicing using full length transcriptome analysis results. In this study, we performed a combined analysis with long-read RNA sequencing (Iso-Seq) and DNA methylation data obtained from the same regions in the same samples to examine the effect of DNA methylation on alternative splicing.

## Results

### Overview of the full-length transcriptome of three brain regions

To characterize the profiles of full-length isoforms in different human brain regions, we extracted RNA from cerebellum, hypothalamus, and temporal cortex, and performed Iso-Seq using PacBio’s Sequel IIe system (Fig. S1a). QC results for raw sequencing data are described in Table S1. The percentage of zero-mode waveguides (ZMWs) that met all of the criteria of the Sequel IIe system exceeded 80% in all samples (Table S1). For each sample, we generated 225–401 Gb of raw sequencing data and 7–14 Gb of HiFi data satisfying >3 full-pass sub- reads and QV ≥20 (Table S2). Reads supported by at least two full length non-concatemer (FLNC) reads with accuracy >99% were mapped to the human genome (hg38) and unique isoforms were clustered using IsoSeq v3 software (Fig. S1b). We checked the number of FL, FLNC and FLNC with polyA reads and confirmed that all sample reads were ≥2,000,000 and that the read number was not related to the RNA quality number (RQN) (Fig. S2). Further QC and filtering of the data using SQANTI3 yielded an average of 27,643 isoforms per sample and 66,860 non-redundant isoforms across all samples (Fig.1a, Table S3). These isoforms were annotated to 14,382 known and 52 novel genes (Fig.1b). When compared by brain region, the number of isoforms was significantly lower in the cerebellum than in the hypothalamus and temporal cortex (Ce vs. Hy: *P* = 8.04E-3, Ce vs. Tc: *P* = 2.74E-4) (Fig.1c). The proportion of novel isoforms commonly expressed in the three regions was 42.9%, whereas the proportion of novel isoforms expressed at only one or two of the regions was higher (61.2–82.2%) (Fig.1a).

We then compared the detected isoforms to those in the comprehensive human gene annotation database GENCODE v.41 and categorized them into the following eight categories: full splice match (FSM), incomplete splice match (ISM), novel not in catalog (NNC), novel in catalog (NIC), fusion, genic, antisense, and intergenic (Fig. 1d). Since the number of fusion, genic, antisense, and intergenic isoforms was very small, we focused mainly on FSM, ISM, NIC, and NNC isoforms. In the data of this study, approximately 57.3% of the isoforms were FSM, 27.9% were NIC isoforms and 8.5% were NNC isoforms. ISM did not account for a large proportion (5.8%) of the identified isoforms (Fig. 1d), which may be due to the strict filtering applied to ISM isoforms, as these are difficult to distinguish from degradation products and many ISM isoforms were considered to be artifacts. Indeed, QC of SQANTI3 with these final isoforms showed that only ISM isoforms had a “Coverage Cage” of around 30–60%, which was lower than that observed for the other categories (Fig. S3). “Coverage Cage” is one of the indicators used for good quality isoforms and a measure of whether a peak is detected by the CAGE method within 50 bp. All good quality indicators for the other categories generally exceeded 75%, which assured the quality of the filtered isoform (Fig. S3).

**Fig. 1.**
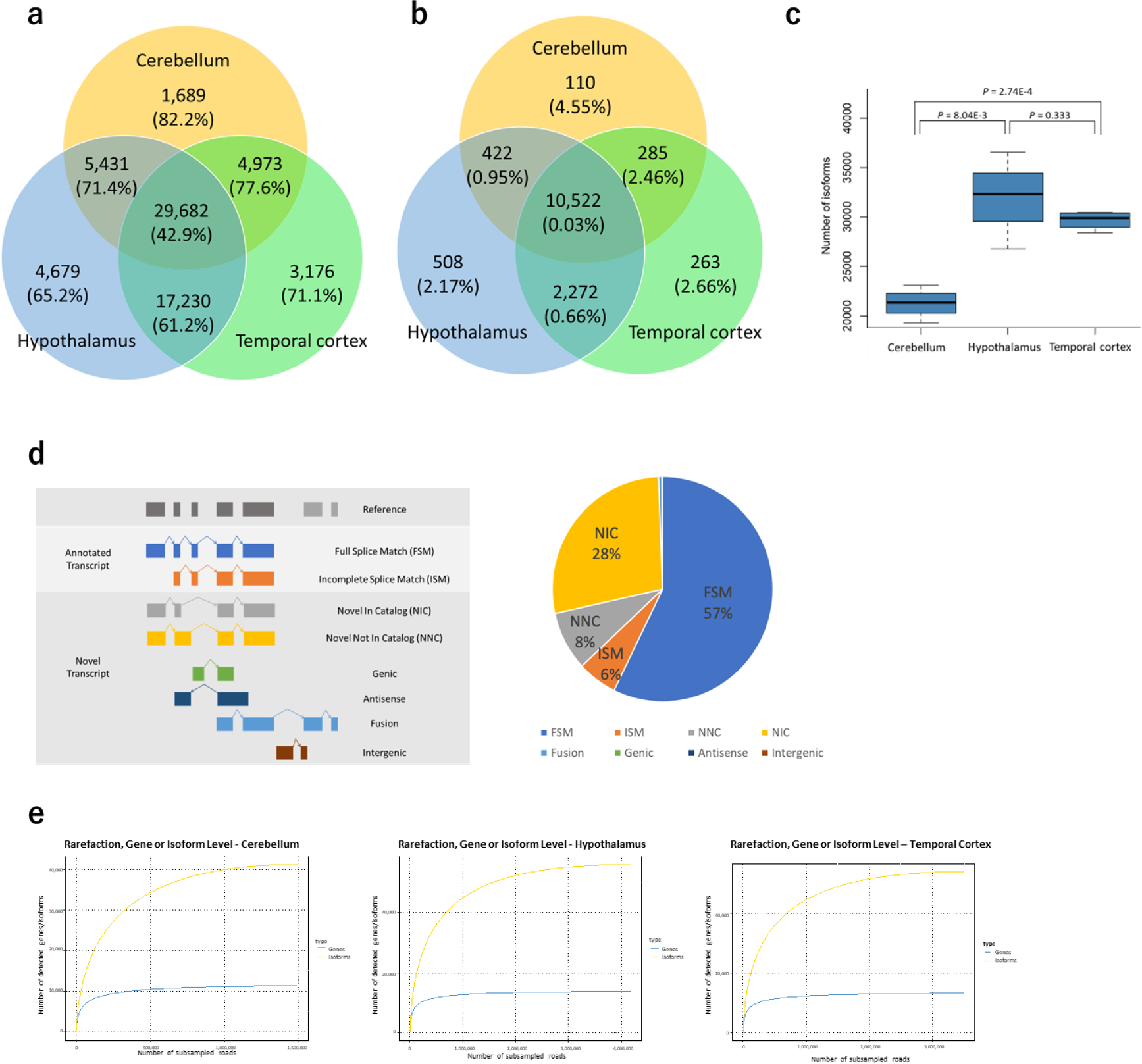
**Isoforms detected in human cerebellum, hypothalamus, and temporal cortex using Iso-seq**. **a** Number of isoforms detected in cerebellum, hypothalamus, and temporal cortex and percentage of novel isoforms indicated in brackets**. b** Number of genes detected in the three brain regions and percentage of novel genes indicated in brackets. **c** Number of detected isoforms by brain region. **d** Definition of isoform categories and breakdown of detected isoforms. FSM: isoform with all splice junctions matching the reference, ISM: isoform with fewer exons than existing splice junctions, but matching the known splice junctions, NIC: isoform with combinations of known donor-acceptor sites, NNC: isoform with at least one novel donor-acceptor site, Genic: within the known genes, Antisense: reverse stranding of known gene regions, Fusion: fusion gene, and Intergenic: outside of known genetic regions. **e** Rarefaction curves of genes and transcripts detected in each brain region.

Next, we performed rarefaction analysis to examine whether isoforms and genes were adequately detected in the sequencing analyses performed in this study. The analysis performed for each gene and isoform showed that the gene rarefaction curves were well saturated in all three brain regions (Fig. 1e). Since the isoform rarefaction curves were also mostly saturated in all three brain regions, but the degree of saturation was not as high as that of the genes, it is possible that some new isoforms could be identified if the depth of sequencing is increased further (Fig. 1e). When analyzed by isoform category, the curves were generally saturated in all regions and in all categories (Fig. S4).

Further, cluster analysis and PCA were performed to examine the relationship between the isoform profiles of each sample. In terms of transcription profiles, the cerebellum differed from the hypothalamus and the temporal cortex, while the hypothalamus and the temporal cortex did not differ much (Fig. S5a, b). Since the transcriptome profile was expected to be markedly affected by the proportion of cells constituting the sample, we estimated the proportion of cells using MeDeCom, a reference-free estimation method [38], using data from a genome-wide DNA methylation analysis that was performed on the same samples used for the Iso-Seq analysis. The estimation results were not homogeneous across samples in the hypothalamus, and some samples were found to have a similar cellular composition to the temporal cortex (Fig. S5c).

### Comparison of isoform features between categories and brain regions

We investigated the characteristics of isoforms and compared them between isoform categories and brain regions. First, we examined how many samples expressed isoforms in each of the eight categories. Since the isoforms that were expressed in only one sample were removed in order to consider more plausible isoforms, those in the analysis were expressed in at least two samples. Compared to novel isoforms, the proportion of FSM isoforms was higher in many samples (average 6.51), and the number of samples detected decreased with increasing degree of novelty, ISM (average 6.12, *P* = 4.80E-16), NIC (average 4.05, *P* <2.2E-16), and NNC (average 3.29, *P* <2.2E-16) (Fig. 2a). As for brain regions, isoforms expressed in all three regions were most abundant in ISM, followed by FSM; compared to FSM, NIC and NNC had significantly higher percentages of isoforms expressed in only one or two regions (NIC: *P* < 2.2e-16, odds ratio (OR) = 2.79, NNC: *P* < 2.2e-16, OR = 4.97) (Fig. 2b). Expression levels (transcripts per million (TPM)) tended to be higher for FSM (Fig. S6a) and for isoforms that were expressed in more brain regions (Fig. S6b).

**Fig. 2.**
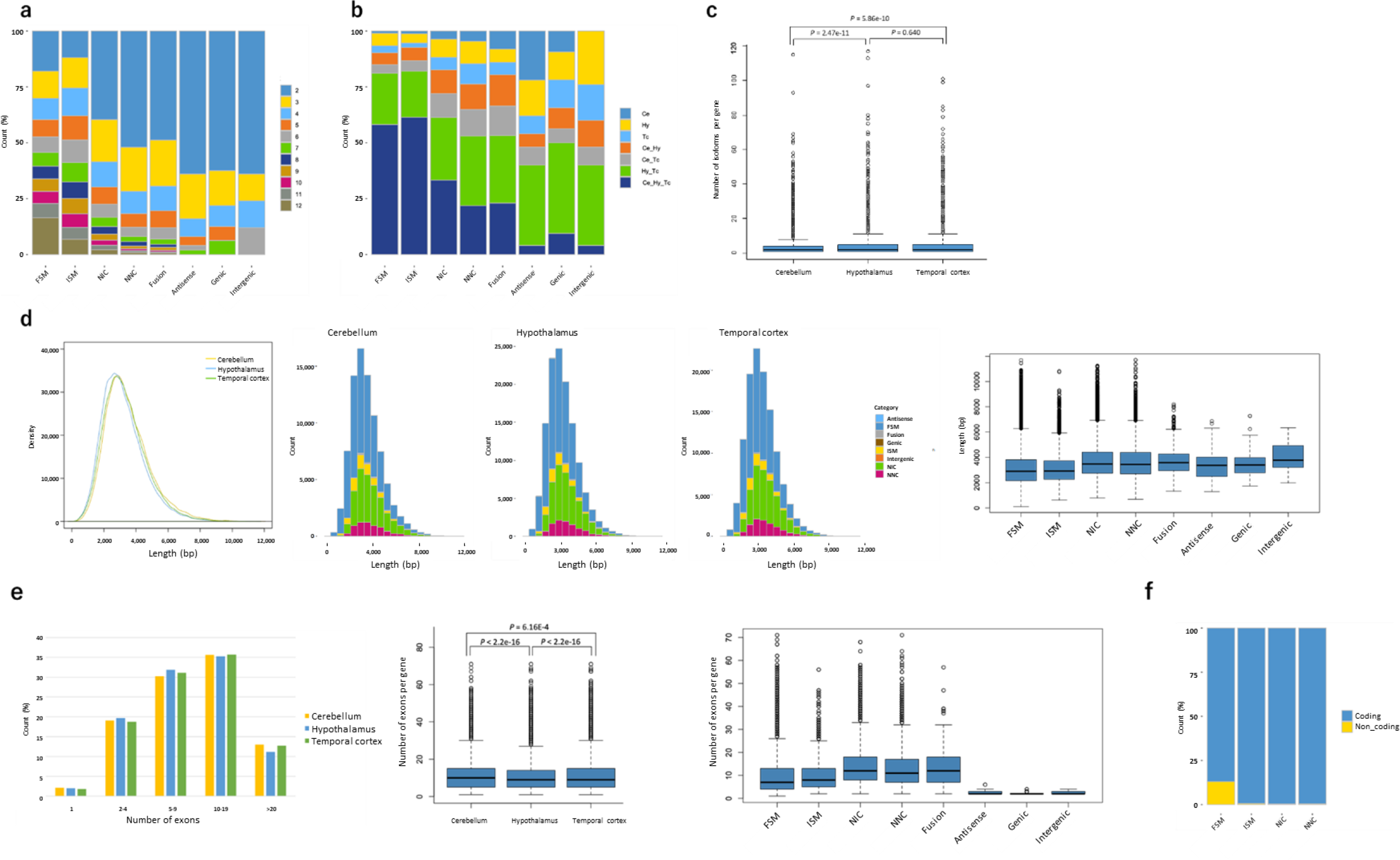
Overview of the detected transcripts by category or brain region. a. Distribution of the number of samples in which the detected isoforms were found. Only isoforms that were expressed in at least two samples were included in subsequent analyses. **b** Distribution of the brain regions in which the detected isoforms were found. Ce, cerebellum, Hy, hypothalamus, Tc, temporal cortex. **c** Number of isoforms per gene in each brain region. **d** Distribution of the read-lengths of all detected isoforms by brain region. In the major four categories, NIC and NNC showed a tendency to have longer lengths. **e** Distribution of the number of exons per gene. The number of exons per gene is higher in the cerebellum and lower in the hypothalamus. Novel isoforms tend to have more exons compared to known isoforms based on the category classification. **f** Percentage of estimated protein-coding isoforms.

Next, the number of isoforms per gene was examined and an average of 2.74 isoforms were identified (Fig. 2c). When compared by region, the number of isoforms per gene was significantly lower in the cerebellum than in the other regions (Ce vs. Hy: *P* = 2.47E-11, Ce vs. Tc: *P* = 5.86E-10) (Fig. 2c). We then examined the distribution of isoform lengths. Isoform read lengths peaked at around 3 kb and tended to be slightly shorter in the hypothalamus (Fig. 2d, Fig. S7a). When examining the length of each isoform by category, novel isoforms except ISM were longer in all brain regions (*P* <2.2E-16), as reported in previous studies [19, 39] (Fig. 2d, Fig. S7b, c). When examining the number of exons per gene in each brain region, the number was higher in the cerebellum (average 11.1 per gene) and significantly lower in the hypothalamus (average 10.6 per gene, *P* <2.2E-16) (Fig. 2e). When compared by category, the distribution of the number of exons per gene was significantly different between FSM (average 9.32 per gene) and novel isoforms (Fig. 2e), especially NIC (average 13.5 per gene, *P* <2.2E-16) and NNC (average 12.8 per gene, *P* <2.2E-16), which was consistent between regions (Fig. S8). For FSM, ISM, NIC, and NNC, which were not filtered based on the condition of ’protein-coding’ during QC with SQANTI3, the proportion of protein-coding isoforms was examined. The results showed that 13.0% of the FSMs were non-coding, while non-coding isoforms of the other novel isoforms were ∼ 0.5%, indicating that most of novel isoforms were predicted to be protein coding (Fig. 2f).

To validate the novel isoforms detected in this study, we then utilized proteome analysis data of postmortem human brain cortex reported in a previous study [40]. Liquid chromatography coupled to tandem mass spectrometry (LC-MS/MS) results were used to detect FDR < 0.01 proteins with both novel isoform sequences and known protein sequences from GENCODE as a reference. Among the amino acid sequences used to detect novel isoforms, those were not detected when using known protein sequences as a reference and not identical to any known sequences were searched. We identified 37 amino acid sequences derived from 34 proteins (Table S4), which is evidence of translated novel isoforms. Alternative splicing events such as exon skipping (Fig. S9a) and intron retention (Fig. S9b) of NIC, and novel splice junction of NNC (Fig. S9c) were validated by the identified amino acid sequences.

### Comparison of gene and isoforms between brain regions

We first examined genes that were highly expressed throughout the brain. The TPM for each gene is shown in Table S5. There were 201 genes with TPM >200, of which Myelin Basic Protein (*MBP*) was the most highly expressed gene throughout the brain, followed by Glial Fibrillary Acidic Protein (*GFAP*). Pathway analysis using these genes detected numerous pathways for nervous and brain development and neurogenesis (Fig. S10a, Table S6). Similar trends were observed when brain regions were examined separately. Most of the pathways detected in analyses of each brain region overlapped with pathways that were detected in analyses of the all three regions, and many of the genes that were highly expressed in each region were related to brain and nervous system pathways (Table S7, S8, S9).

Next, we searched for genes that were only expressed in each respective region. Very few genes were expressed in only one of the three brain regions, and only 42 genes met our criteria for the cerebellum (Table S10), 25 in the hypothalamus (Table S11), and 44 in the temporal cortex (Table S12). Pathway analysis using these region-specific genes detected no significant pathways in the analyses of the cerebellum and hypothalamus, while synaptic signaling-related pathways were detected in the analysis of the temporal cortex (Table S13).

Since Iso-Seq was considered to be less suitable for differential expression analysis than short read analysis [41], we searched for genes that were differentially expressed in each brain region based not only on statistical significance, but also on the strict criterion of fold change (FC) >5. The number of genes upregulated in the cerebellum according to our criteria was 58 (Table S14), including Zinc-finger protein of the cerebellum (ZIC) family members *ZIC1*, *ZIC3*, and *ZIC4*, and pathway analysis showed that the top pathways included “neuronal pathways”, “ion transport”, and “brain development” (Fig. S10b, Table S17). Similar pathways involved in neurogenesis were detected in the analysis using 77 upregulated genes in the temporal cortex (Fig. S10d, Table S16, Table S19). In the hypothalamus, on the other hand, 53 genes with elevated expression were detected (Table S15), and many of these genes were involved in the “immune response” and “cell migration” pathways (Fig. S10c, Table S18). As for the site-specific down-regulated genes, only 25 were detected in the hypothalamus and 4 in the temporal cortex, whereas 314 were detected in the cerebellum (Table S14, 15, 16). Pathway analysis showed that many of the cerebellum-dependent genes with decreased expression were involved in “actin filament-based process”, “anatomical structure” and “cellular component size” (Fig. S10e, Table S17).

We then focused on genes with different numbers of isoforms per gene. First, we searched for genes expressing more than 20 isoforms per gene in the whole brain, and 488 genes were detected (Table S20). Pathway analysis using these genes identified “macromolecule and protein localization” and “cell projection organization” pathways in addition to brain-specific pathways such as “neurogenesis” (Fig. S11a, Table S21). To investigate whether this trend was brain-specific, we used long-read data from the GTEx [20] to detect genes with high isoform counts in cells and tissues other than the brain and performed a pathway analysis using these genes. The results showed that five of the top 10 pathways, including “macromolecule and protein localization”, were detected in the analysis of tissues other than the brain, and that the genes involved in these pathways generally expressed numerous isoforms (Fig. S11a, Table S21). Next, we searched for genes with a particularly high number of isoforms per gene in the regions of interest compared to other regions, and found 34 genes in the cerebellum, 136 genes in the hypothalamus, and 77 genes in the temporal cortex (Table S22, 23, 24). No significant pathways were detected in the analysis of genes that had a higher number of isoforms per gene in the cerebellum, but these genes included *ZIC3* and *ZIC4*, which were also detected as a cerebellum-dependent expressed genes (Table S22). Pathway analyses of the hypothalamus and temporal cortex detected such pathways as “membrane and cytoskeleton organization” and “immune response” in the hypothalamus and “neuron differentiation” and “cell projection organization” in the temporal cortex, respectively (Table S25, 26, Fig. S11b, c). The correlation between the number of isoforms per gene and expression levels was not strong (r^2^ = 0.127, Fig. S12); however, pathways related to the immune response in the hypothalamus and neurogenesis in the temporal cortex were detected, not only for genes that were highly expressed in a region-dependent manner, but also for genes with a high number of isoforms.

### Genes with different major isoforms in different brain regions

Next, we searched for genes with different major isoforms in different regions. For each sample, the ratio (%) of the expression of individual isoforms to total gene expression was calculated, and the ratios and their ranks were compared among regions. To determine whether major isoforms were differed among regions, we used both parametric and nonparametric tests to search for isoforms with ratios that differed by more than 20% (*P* <0.05), and with ranks of expression that differed between regions. As a result, 193 isoforms of 145 genes were identified in the cerebellum vs. the hypothalamus, 147 isoforms from 114 genes were identified in the cerebellum vs. the temporal cortex, and 55 isoforms from 47 genes were identified in the hypothalamus vs. the temporal cortex (Table 1, Table S27, 28, 29). Among the genes for which isoforms were detected in all three comparisons, Growth Arrest Specific 7 (*GAS7*) was the gene with the lowest P value (Table S27, 28, 29). The transcripts ENST00000580865.5 and ENST00000437099.6 were predominantly expressed in the cerebellum and temporal cortex, respectively (Fig. 3a). In the hypothalamus, both ENST00000585266.5 and a novel isoform, TALONT000412184, were expressed (Fig. 3a). In a previous study using western blotting in mouse brain, different isoforms of *GAS7* that were predominantly expressed in the cerebellum and cerebral cortex were reported [42]. Therefore, we compared the predicted amino acid sequence encoded by the isoforms detected in this study with those of mice. As a result, the longer isoform, Gas7, expressed in the mouse cerebrum showed 97.33% identity with ENST00000585266.5 detected in the hypothalamus and 97.35% identity with ENST00000437099.6 detected in the temporal cortex. On the other hand, the shorter isoform, Gas7-cb, expressed in the mouse cerebellum showed 97.9% identity with ENST00000580865.5 detected in the cerebellum (Fig. 3b).

**Fig. 3.**
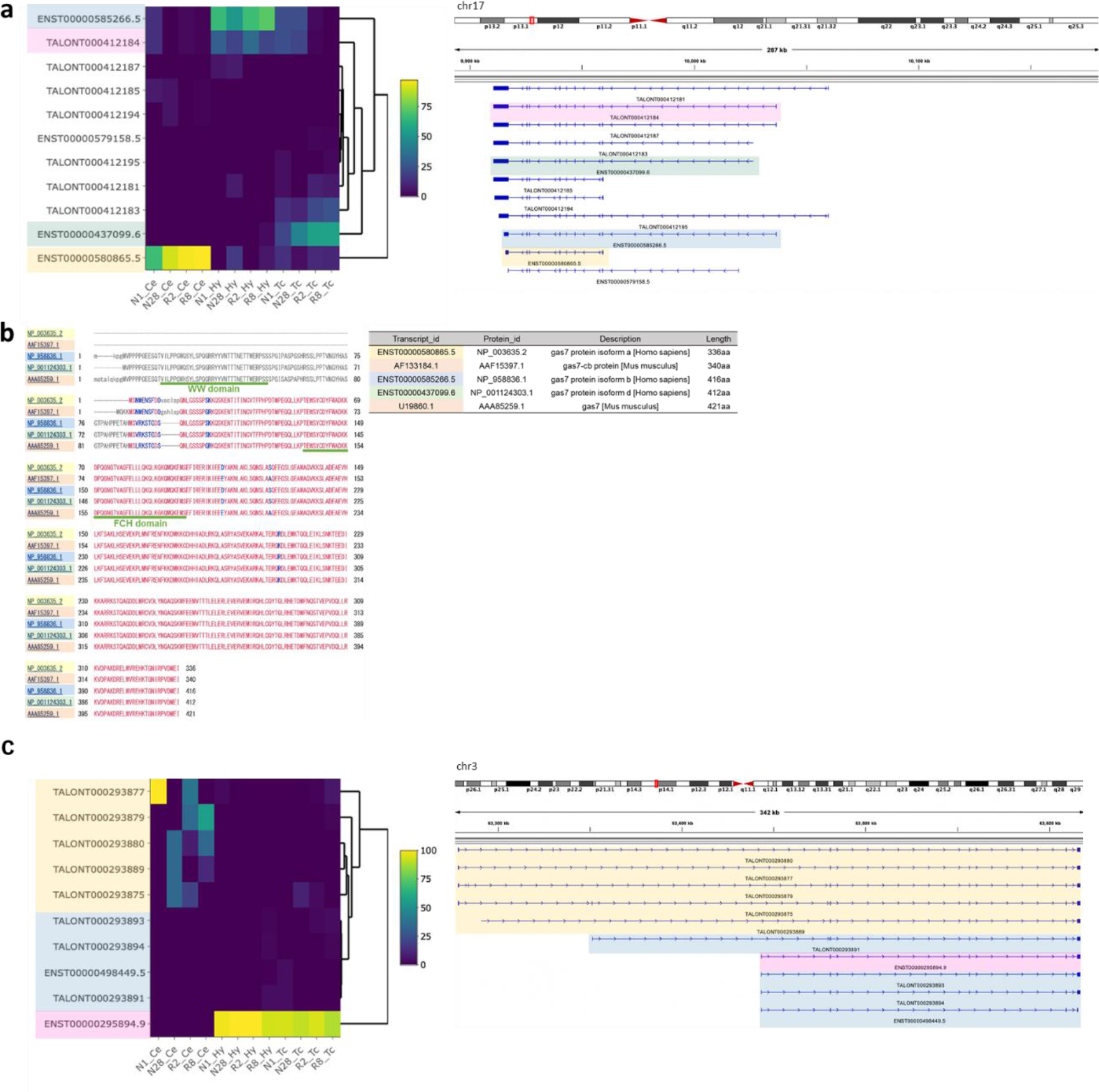
Examples of a gene that uses different major isoforms depending on the brain region. a. *GAS7* is an example of a gene with different major isoforms at each region. The heatmap represents the proportion of expression levels of each isoform relative to the overall expression levels of the gene. The colors on the transcript IDs correspond to the colors of the transcript structures displayed in IGV. **b** Comparison of predicted amino acid sequences between transcripts detected in this study and those reported to be expressed in the mouse cerebellum (AF133184.1/AAF15397.1) and cerebral cortex (U19860.1/AAA85259.1). **c** An example of *SYNPR* found to have isoforms that are the most significantly differentially expressed between Ce and Hy, and was also the second most significant when comparing Ce and Tc.

**Table 1.**
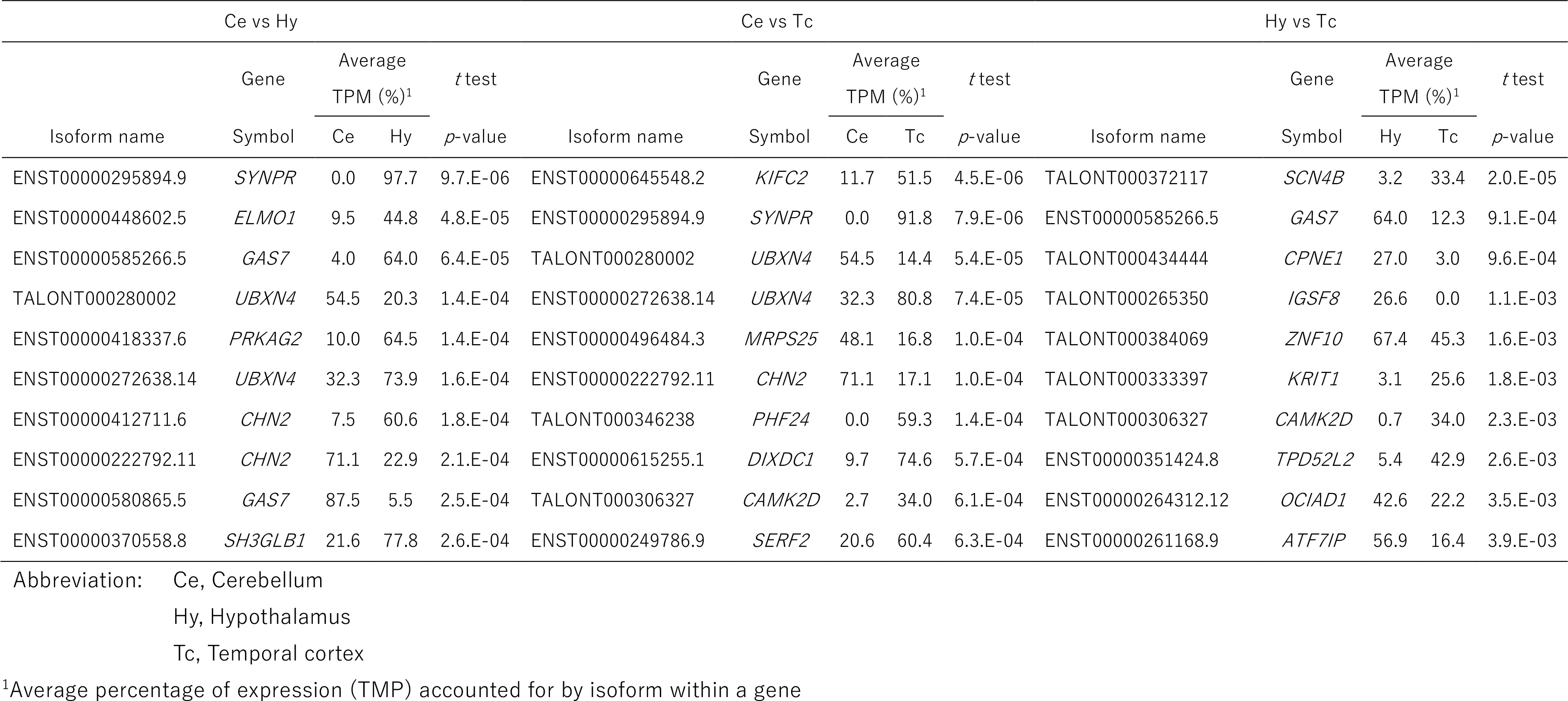
Top 10 isoforms for which the proportion of gene expression differs depending on the brain region

The isoform with the lowest P value in the cerebellar and hypothalamic comparison was that of Synaptoporin (*SYNPR*), which was one of the top two isoforms in the cerebellar and temporal cortex analysis (Table S27, 28). ENST00000295894.9 was expressed in the hypothalamus and the temporal cortex, whereas the cerebellum expressed five new, longer isoforms with transcription start sites (TSS) more than 150 kb upstream of ENST00000295894.9 (Fig. 3c).

By analyzing the genes that expressed different major isoforms between the cerebellum and hypothalamus, we identified pathways related to “cell projection organization”, “cellular component organization/biogenesis” and “actin filament-based process” (Fig. 4a, Table S30). Similarly, when we analyzed genes with different major isoforms between the cerebellum and temporal cortex, we found their involvement in “macromolecule and protein localization” (Fig. 4b, Table S31). Finally, we identified “actin filament-based process” as a pathway associated with genes exhibiting different major isoforms between the temporal cortex and hypothalamus (Fig. 4c, Table S32). Pathways such as “actin filament-based process”, “cell projection organization”, and “cellular component organization/biogenesis” were identified in multiple comparisons due to the influence of genes such as *GAS7*, which was detected in all three comparisons, or specific isoforms expressed at only one site and therefore detected in two comparisons. In addition, although not included in the top 10, pathways such as “neuron differentiation” and “neurogenesis” were also identified as significant pathways in the cerebellum vs. the hypothalamus comparison, indicating that a certain number of genes which highly express different isoforms in different brain regions were also responsible for nervous system functions (Table S30, 32).

**Fig. 4.**
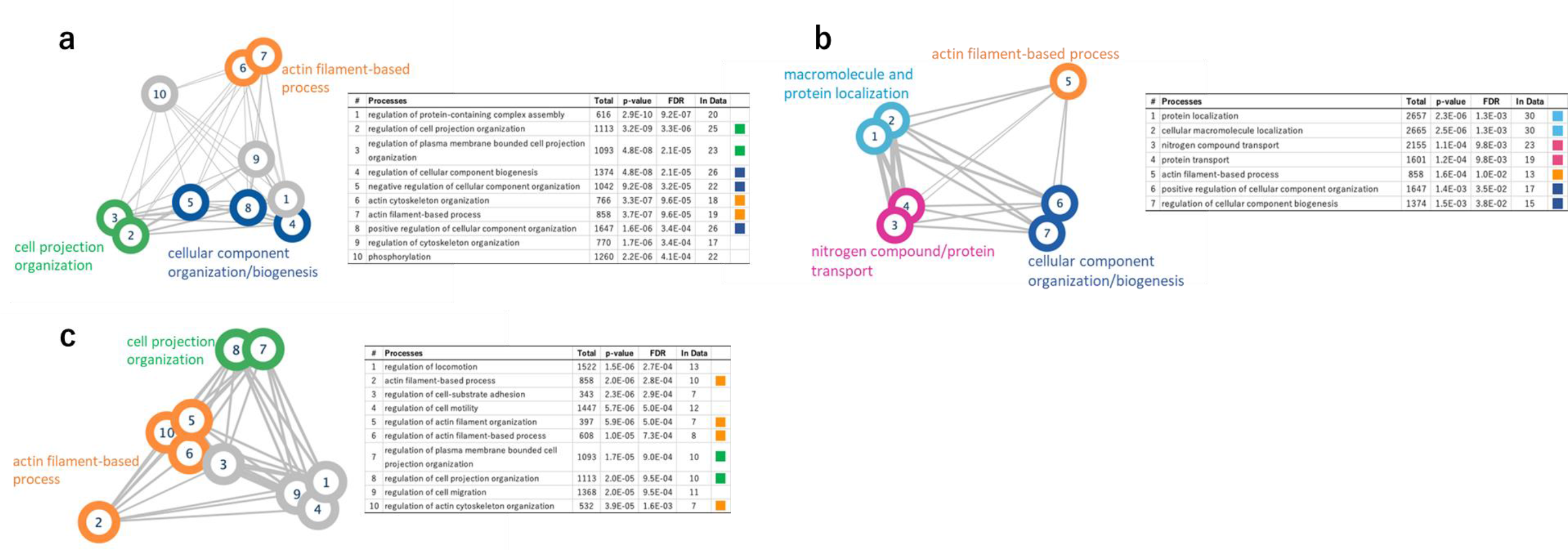
Pathway analysis of genes expressing different major isoforms in different regions of the brain. When more than 10 pathways are detected, the top 10 pathways with the smallest false discovery rate (FDR) are shown. Edge thickness is proportional to the number of related genes shared among pathways. **a** Pathway analysis results for genes with different major isoforms expressed in the cerebellum and hypothalamus. **b** Pathway analysis results for genes with different major isoforms expressed in the cerebellum and temporal cortex. **c** Pathway analysis results for genes with different major isoforms expressed in the hypothalamus and temporal cortex.

### Integrative analysis of transcriptome and methylation data

We identified several genes that expressed different isoforms in different brain regions. The samples used in this study were comprised of different brain regions of the same samples and there were no differences in genomic sequences (e.g., SNPs) among regions, and the effects of environmental factors were also expected to be similar between regions. Therefore, considering the possibility that epigenetics may play an important role as a mechanism for the expression of different isoforms within a gene, we focused on DNA methylation. Using data from a previous study [43] in which we performed array-based whole-genome methylation analyses using DNA extracted from the same samples used in this study, we first searched for differentially methylated positions (DMPs) between regions and identified 66,908, 32,841 and 8,752 DMPs in comparisons between the cerebellum and the hypothalamus, the cerebellum and the temporal cortex, and the hypothalamus and the temporal cortex, respectively. We examined whether these DMPs were abundant in the vicinity of genes with different major isoforms in the different regions. The results showed that DMPs were significantly more abundant around genes with different major isoforms between the regions than around all of the expressed genes in the brain in this study (Ce vs Hy; *P* = 3.87E-07, Ce vs Tc; *P* = 4.48E-05, Hy vs Tc; *P* = 4.51E-02) (Table 2). When we examined the areas where these DMPs were present, we found that DMPs were more common in the region 200-1500 bp upstream of TSS, 5’ UTR and first exon (Table 3). We also examined the relationships between DMPs and genomic region features and CpG islands, and found that DMPs were abundant in the enhancer regions and transcription factor binding sites; however, the relationship between DMPs and CpG islands was not strong (Table 3).

**Table 2.**
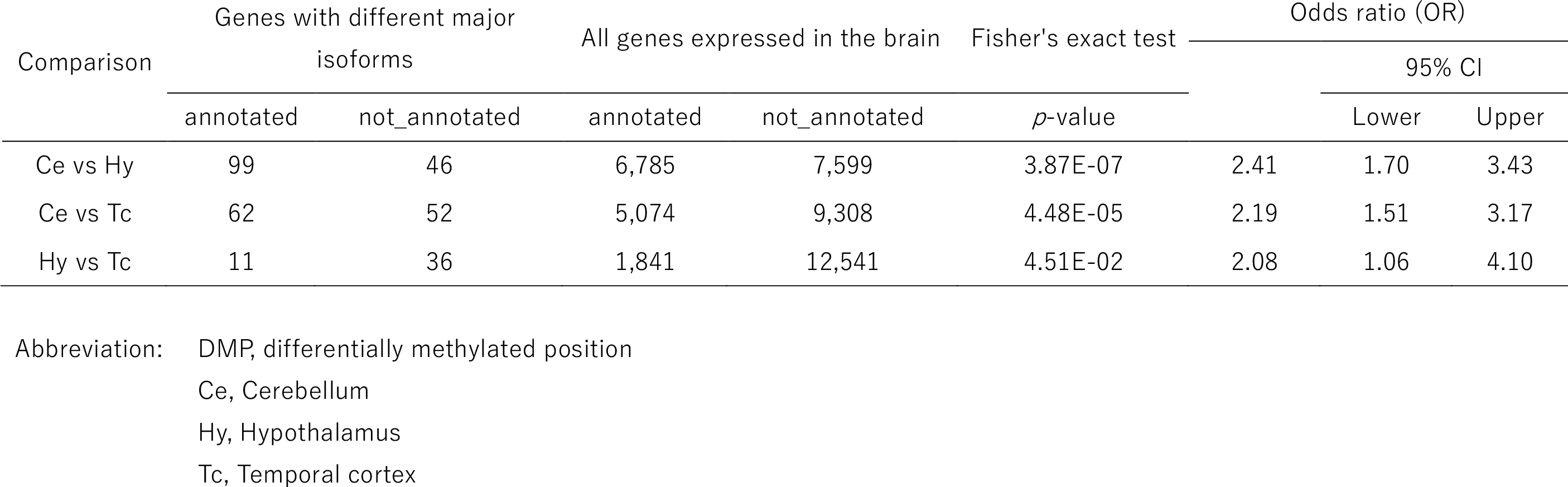
DMPs around genes with different major isoforms expressed in each region

**Table 3.**
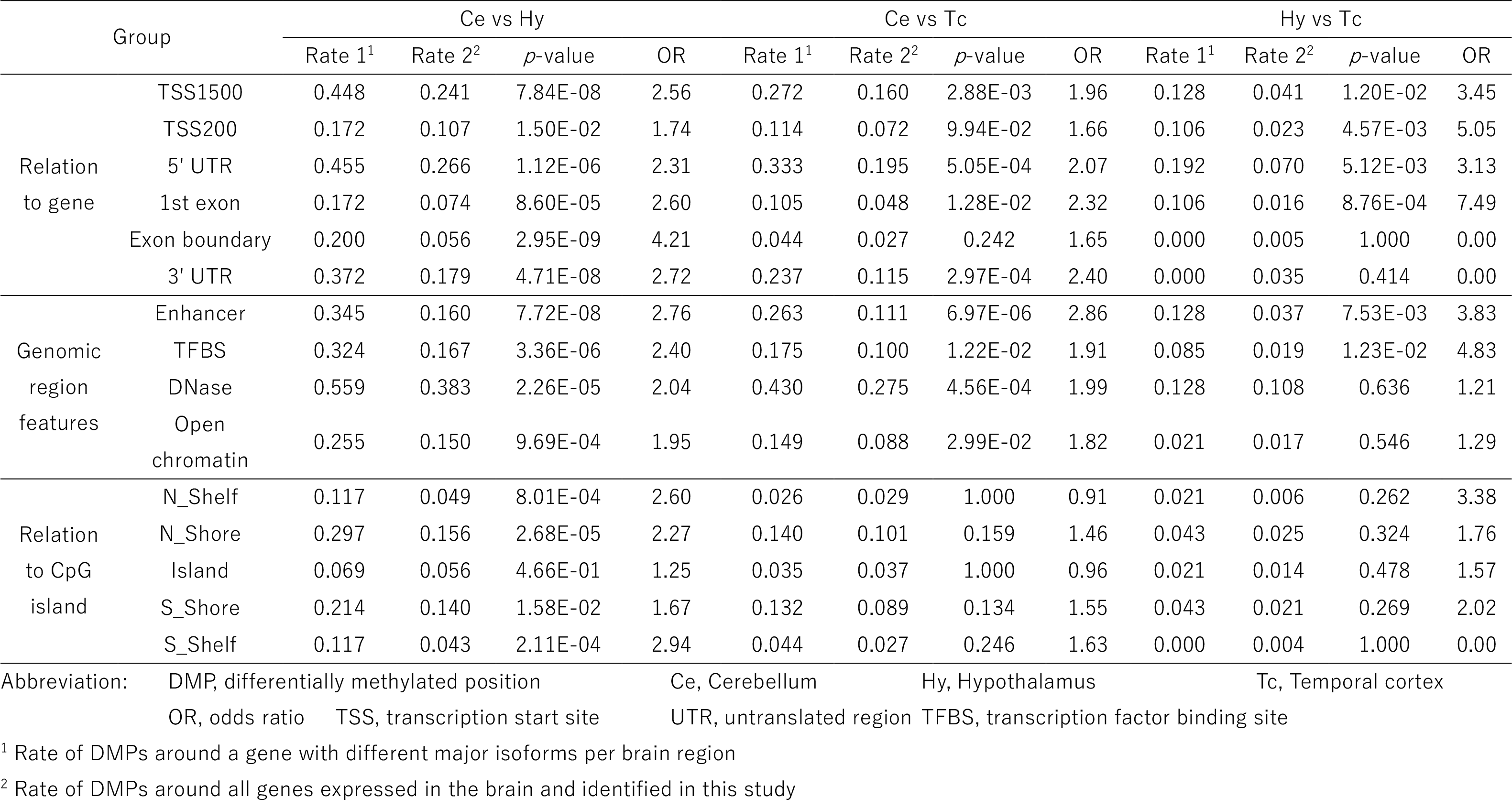
Characteristics of areas with DMPs around genes with different major isoforms in each region.

Given the existence of many DMPs around genes with different major isoforms between regions, we examined methylation around *GAS7* and found that there were many DMPs, especially around TSSs where isoforms of different lengths were expressed (Fig. 5a). In order to examine methylation associations in more detail for the differences in isoform structures, we defined the pair of isoforms to compare (Fig. 5b). Besides, the structural differences between isoforms were categorized into isoforms with different first and/or last exons, the same exon with different 5’ and/or 3’ splice sites, intron retention, exon skipping, cassette exons or other complicated exon structures (Fig. S13). In each case, the DMPs of the associated regions were explored. To compare the degree of DMP accumulation, the same analysis was performed for control isoforms, the expression patterns of which did not differ among regions. The distribution of the pattern of differences in isoform structure did not differ markedly between genes with different major isoforms in the different regions and those of the control isoforms (Fig. 5c). Next, we examined the DMPs in the vicinity of related regions for isoform structural differences. Although the number of identified structural patterns, such as cassette exons and exon skipping were too small to evaluate, especially in the hypothalamus vs the temporal cortex comparisons (Table S33), in the comparisons of the cerebellum vs. the hypothalamus and the cerebellum vs. the temporal cortex, we found that there were significantly more DMPs within regions 1 kb downstream from the TSSs of the isoforms that were differently expressed between regions and had different first exons (Ce vs Hy; *P* = 3.47E-04, Ce vs Tc; *P* = 4.48E-08) (Fig. 5d, Table S34). There were also more DMPs of nominal significance in areas 1 kb upstream of the isoform TSSs that varied in expression between regions, but the degree of association was not as strong as in 1 kb downstream area (Ce vs Hy; *P* = 5.61E-03, Ce vs Tc; *P* = 0.023) (Fig. 5d, Table S34). Genes with DMPs within 1 kb downstream of the TSSs of isoforms with different first exons are shown in Table S35.

**Fig. 5.**
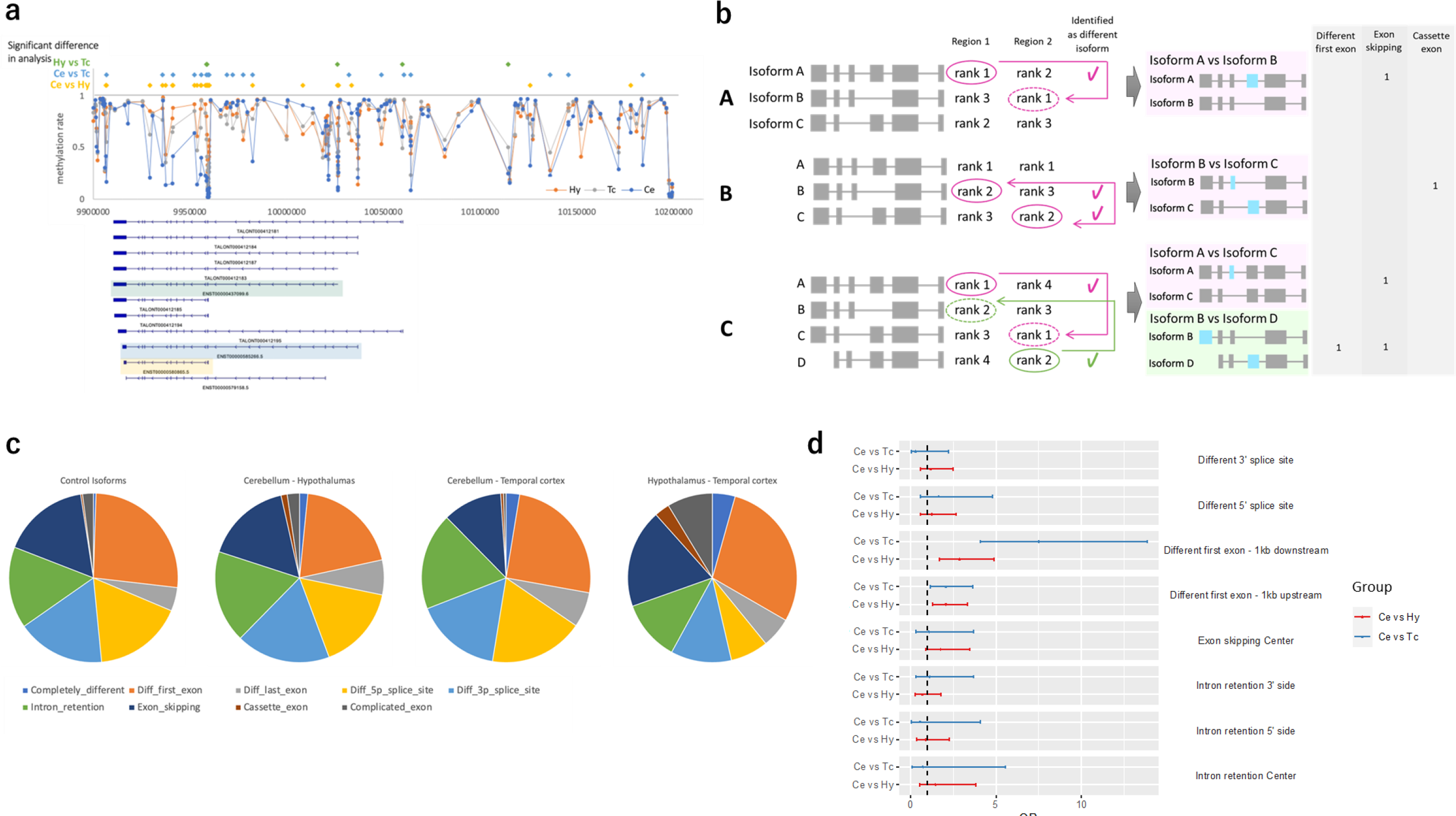
Relationship between genes that have different isoforms in different brain regions and DNA methylation status. a. The DNA methylation status in the vicinity of *GAS7* in each brain region. Numerous differentially methylated positions (DMPs) exist between brain regions in the vicinity of the transcription start site of ENST00000580865.5, which is an isoform that is expressed in the cerebellum. **b** Strategies for selecting isoforms to compare. If a difference in the proportion of expression of a certain isoform between region 1 and region 2 is detected, the rank of the isoform in terms of expression level within the gene is compared between the regions, and the lower rank is identified. The isoform corresponding to that rank in the other region is used for a comparison. **A.** If isoform A with the highest expression level (rank 1) in region 1 is detected, then the isoform that ranks first in region 2 is selected as the comparison target. **B.** If isoform A is ranked first in both regions and isoform B with the second highest expression (rank 2) in region 1 is detected, then isoform C with a rank of 2 in region 2 is selected as the comparative isoform. In some cases, isoform C is also detected as having a significantly different expression ratio between regions. **C.** Isoforms A and D have been detected, and isoforms corresponding to the lower ranks, rank 1 and rank 2, are selected as comparative isoforms. **c** Proportion of structural differences in isoforms with different expression ratios between brain regions. The control isoform was defined as the isoform with one of the top two most highly expressed isoforms in at least one region of the brain, and was consistent across all regions without any significant differences in expression levels between them. **d** Forest plot of odds ratios (ORs) in the test for whether DMPs are more common in areas associated with the structure of isoforms that are expressed differentially between brain regions.

## Discussion

In this study, we conducted a full-length transcriptome analysis of mRNA extracted from postmortem human brains using long-read sequencing. Long-read sequencing using human brain samples is very rare, and to the best of our knowledge, this is the first study to use samples from the hypothalamus. By analyzing mRNA using long-read sequencing, it is possible to read the entire length of most transcripts at once, allowing for more accurate identification of isoforms compared to short-read RNA-seq-based analyses. As a result, approximately 40% of the isoforms identified in this study were novel. NIC was the most common new isoform identified at 65%, followed by NNC and ISM. The number of ISM isoforms may have been overly reduced because they were filtered strictly using a machine learning filter that was not applied to other isoforms. The reason only ISM isoforms were screened in this way is because ISM isoforms are either likely to contain RNA degradation products or cDNA that were not successfully synthesized experimentally. The novel isoforms NIC and NNC, which accounted for most of the identified novel isoforms, were longer and contained more exons compared to the existing isoforms, which was consistent with the results of previous studies [19, 39]. Most of the new isoforms were presumed to be protein coding according to the SQANTI3 prediction, suggesting that they may have some function in the brain. The detection of peptides derived from the novel isoforms by LC-MS/MS of the human postmortem brain samples further supports the possibility that the novel isoforms are translated and functional.

We focused on genes and isoforms that are differentially expressed in different regions of the brain, i.e., the cerebellum, the hypothalamus, and the temporal cortex. The samples used in this study were all postmortem brains from men in their 50s or older, and the comparison of different regions of the same samples minimized the effects of genetic variations and environmental factors on the analysis. When we compared gene expression levels in different regions of the brain, we found that many of the genes involved in neural pathways were highly expressed throughout the brain, and this trend was also observed in each brain region. Additionally, we found that the number of genes that were only expressed in one of these regions was very small. This is likely because the brain samples used in this study were comprised of a mixture of cell types. If we were to perform our analysis on single cells, then the likelihood of identifying more cell-specific genes would be increased. This assumption is supported by the methylation data of our samples, which showed that putative cell types are shared between different brain regions. We did find some genes that showed differences in expression levels between different regions of the brain. As long-read RNA-seq is less quantitative compared to short-read sequencing methods, we identified these genes using a particularly stringent criterion of FC>5. In the cerebellum, we found that many genes involved in “neurogenesis”, as well as in “head, brain, and central nervous system development” pathways, showed higher expression levels compared to other regions of the brain. Of particular interest, the *ZIC* family genes including *ZIC1*, *ZIC3*, and *ZIC4* were identified. *ZIC* genes encode a group of evolutionarily conserved proteins with zinc finger motifs that are known to regulate cell differentiation and are involved in neural and early development [44–46]. The cerebellum is derived from the dorsal hindbrain, where the *ZIC* genes are strongly expressed, and in adults, *ZIC* is expressed only in cerebellar granule cells and a few other nuclei and is therefore commonly referred to as the “zinc finger protein of the cerebellum” [47]. *ZIC3* and *ZIC4* were also detected as genes with a higher number of isoforms in the cerebellum than in other regions in this study, again suggesting that these genes play an important role in the cerebellum.

Analysis of genes that were specifically upregulated in the temporal cortex also revealed pathways related to neurogenesis, as in the cerebellum. However, in the hypothalamus, immune response pathways were detected exclusively. The reason why the expression patterns of genes involved in immune pathways is detected differently in the hypothalamus is not certain; however, hypotheses include the possibility that the infiltration of immune cells is greater in the hypothalamus than in other brain regions and that immune process and metabolism regulation by cytokines are specific to the hypothalamus [48, 49]. In some regions of the brain, the blood-brain barrier has been modified to form a more permeable barrier to allow certain substances from systemic circulation to access the central nervous system [50]. One such region, the median eminence, is in physical proximity to the hypothalamus, making the hypothalamus more accessible to immune cells and other substances than other brain regions [48, 50]. Our results appear to corroborate the existence of such immune processes in the hypothalamus, but further studies are necessary. Next, we searched for genes that express more than 20 isoforms. Analysis of the entire brain identified numerous genes that are involved in pathways such as anatomical structure and cellular component size, as well as actin filament-based processes. We also performed a similar analysis using data from previous studies that used long-read transcriptome sequencing in tissues other than the brain. Our findings showed that the top 5 pathways were detected in both brain and non-brain tissues, and suggest that the genes involved in these pathways have a high number of expressed isoforms not only in the brain, but also in many other tissues. After searching for genes with a higher number of expressed isoforms in specific regions of the brain and performing pathway analysis, we identified pathways involved in the nervous system and cell projection organization in the cerebellum and temporal cortex. However, in the hypothalamus, in addition to pathways involved in membrane and cytoskeleton organization, immune response pathways were also detected. We then examined the correlation between the number of isoforms and expression levels per gene, and found that the correlation was not strong (r^2^ = 0.127), and that genes with higher expression levels in the hypothalamus did not always have a higher number of isoforms. These findings suggest that, as with genes with high expression levels, genes with a high number of isoforms may also play important functions in pathways such as neurogenesis in the cerebellum and temporal cortex, as well as in pathways of the immune system in the hypothalamus.

We then performed an exploration of genes that express different major isoforms in each brain region, which is the main focus of our research. As a result, the gene *GAS7* was found to show the most significant difference in the major expressed isoforms among the three brain regions. It has been reported that *GAS7* not only has important functions in neurite outgrowth [51, 52], but is also involved in neuronal mitochondrial dynamics and is required for normal neuronal function [53]. In a previous study, the expression of two murine isoforms of GAS7, Gas7 and Gas7- cb, were examined by western blotting [42]. The findings showed that Gas7 is expressed in the cerebral cortex and growth-arrested NIH3T3 fibroblasts, while Gas7-cb is predominantly expressed in the cerebellum. The shorter isoform, Gas7-cb, which consists of 340 amino acids, was found to have an almost identical amino acid sequence to ENST00000580865.5, which was predominantly expressed in the human cerebellum in this study. The longer isoform consisting of 421 amino acids had high sequence identity with those of ENST00000585266.5 and ENST00000437099.6, which were predominantly expressed in the hypothalamus and temporal cortex, respectively. These results support the differences in isoform expression found in our analysis for each brain region and provide support for the reliability of the isoforms identified in our analysis.

Pathway analysis using genes with different major isoforms in the different regions identified pathways such as “actin filament-based process” and “cell projection organization”. The actin cytoskeleton is a fundamental regulator of cell morphology, including that of neuronal cells, and in neurons, actin is concentrated in dendritic spines. Since cell projections are structures such as axons and dendrites in brain cells, these findings suggest the possibility that some of the genes involved in determining the morphology of axons and dendrites in neurons and glial cells employ different isoforms in the different brain regions. The expression profile of guanine-nucleotide-exchange factors that activate calcium/calmodulin-dependent kinases and small GTPase Rac, which are involved in actin reorganization and spine morphogenesis, has been reported to be regionally controlled in the brain [54, 55], supporting the results of this study.

Finally, we considered the possibility that epigenetics may play an important role in the background of the different usage of isoforms in different brain regions, and showed that there were significantly more DMPs in the vicinity of genes with different isoforms in different regions. The methylation sites were particularly abundant in the upstream regions and enhancer of genes, and when the structural differences between isoforms are considered in detail, it became clear that DMPs are particularly abundant in the 1 kb region downstream of the TSS between isoforms with different first exons. DNA methylation is known to affect gene expression, and in general, methylation of the upstream region of a gene is negatively associated with gene expression, while methylation of the gene body is positively correlated with gene expression [33, 56–58]. However, several studies have reported that DNA methylation is not only related to gene expression, but also to alternative splicing [35–37]. In particular, the regulation of exon inclusion is considered to be one mechanism that contributes to alternative splicing [35, 36]. Our results did not show a clear correlation between methylation and the structural differences in isoforms related to exon inclusion, such as cassette exons or exon skipping. However, this may be due to the limited coverage of the methylation probes in the gene regions, which may have prevented the detection of sufficient effects in this study. Further analysis using sequencing-based methylation analysis could reveal a more detailed picture of methylation sites and their associations with alternative splicing.

Our study has several limitations. First, the sample size was small, and the limited number of samples obtained from each brain region made it difficult to accurately evaluate small differences in expression levels, the number of isoforms expressed, and the isoforms that were differentially expressed in each brain region. Matching sample characteristics such as age and sex as much as possible may have compensated for this limitation to some extent, and strict criteria were used to minimize false positives. However, an analysis with more samples would reveal a more detailed profile of isoforms in the brain. Second, some of the hypothalamus-derived RNA used in this study was of poor quality, and it is possible that some of the mRNA included was degraded. In this study, we applied very strict filtering for ISM and excluded many ISMs from the analysis. By using better quality RNA and relaxing the stringency of filtering conditions, it may be possible to detect truly functionally important ISMs more accurately. Third, our brain region-specific analyses were performed on bulk tissue fragments containing a variety of cells, so it is difficult to determine whether the results are specific to the region or to a specific cell type within that region. According to a recent study which analyzed long reads of single cell-derived RNA from mouse brain cells, the expression of region-specific isoforms is often influenced by differences in cell types, although there are rare cases in which the regional differences themselves can have an impact [13]. Based on the composition of the cells inferred from the methylation rates of our samples, cell composition clearly differed between regions, indicating that the region-specificity observed in this study reflects specific features derived from certain cell types. In the future, single-cell sequencing techniques could be used to obtain isoform profiles of individual cells in specific brain regions. Fourth, our data lack proteomic analysis of the same samples, so it is unclear whether the detected isoforms are translated into proteins and whether they play a functional role. Although we estimated open reading frames (ORFs) and confirmed that many of the novel isoforms are presumed to be protein coding, and some of the novel isoforms were supported by the detected peptide sequences by LC-MS/MS analysis of cortical samples from a previous study, the brain region being investigated in the previous study was different from our samples, and thus further experimental validation is preferable using LC-MS/MS data from the same brain region of the same subjects. Finally, our genome-wide methylation data, which we used to investigate the association with alternative splicing, was based on an array-based analysis. While the probe density is high in the upstream region of genes, the density is not high in the gene body, which prevented adequate detection of associations. Sequence-based methylation data with sufficient coverage would be more helpful for more detailed analysis.

## Conclusions

In summary, we analyzed the full-length transcriptomes of the human cerebellum, hypothalamus, and temporal cortex from the same samples and showed that several genes, including *GAS7*, use region-dependent isoforms, and that DNA methylation might be involved in this mechanism. These genes with different major isoforms across regions are particularly involved in dendritic morphology and neuronal pathways in the nervous system, and they are likely to play important roles in their respective regions of the brain. In the future, detailed analysis of isoform expression in the brain regions responsible for neuropsychiatric disorders is considered to be important for elucidating disease mechanisms and for exploring drug targets. We consider that our data will be a valuable resource for such analyses.

## Methods

### Brain samples and RNA extraction

Three regions –– the temporal cortex, hypothalamus, and cerebellum –– were excised from postmortem brain samples from unrelated Japanese individuals (N = 4 × 3). All of the tissues were from male subjects without brain lesions or infections and who died in their 50s or later. The demographic data of these brain samples are summarized in Table S36. All of the samples were autopsied within a postmortem interval of 6 hr and promptly stored at -80°C at the Niigata University Brain Research Institute Brain Bank, Japan.

RNA was extracted using RNeasy^®^ Plus Universal Kit (QIAGEN, Netherlands) after 10–35 mg of tissues was placed in RNAlater^®^-ICE (Thermo Fisher Scientific, MA, USA) for a minimum of 16 hours at -20°C to stabilize the RNA. RNA quality was assessed using a NanoDrop spectrophotometer (Thermo Fisher Scientific) and Femto Pulse System (Agilent Technologies, CA, USA), and RNA quality number (RQN) was calculated according to the Agilent Technical Overview, “Comparison of RIN and RQN for the Agilent 2100 Bioanalyzer and the Fragment Analyzer Systems”; RQN are generally compatible with RNA integrity number (RIN). Samples with RQN greater than 5.0 were used for sequencing.

### Iso-Seq library preparation and single molecule, real-time (SMRT) sequencing

First stand cDNA synthesis and cDNA amplification were performed with a NEBNext Single Cell/Low Input cDNA Synthesis and Amplification Module (New England Biolabs, MA, USA) and Iso-Seq Express Oligo Kit (Pacific Biosciences (PacBio), CA, USA) according to the procedure and checklist (i.e., Iso-Seq™ Express Template Preparation for Sequel^®^ and Sequel II Systems) provided by PacBio. Amplified cDNA samples were purified with ProNex beads (Promega, WI, USA) using the standard workflow and assuming that most transcripts are ∼2 kb. Quantification was performed using Qubit Fluorometer (Thermo Fisher Scientific) and size distribution was confirmed by a Femto Pulse System.

The Iso-Seq library was prepared by going through the steps of DNA damage repair, end repair and A-tailing, overhang adopter ligation, and cleanup using a SMRTbell Express Template Prep Kit 2.0 (PacBio). Sequencing primer was annealed and polymerase binding, complex cleanup, and sample loading was performed with a Sequel II binding kit 2.1 and internal control 1.0 (PacBio) following the manufacturer’s protocol, and sequencing was performed using the Sequel IIe system. Using one SMRTcell per sample, movie runtime was set to 24 hr for each SMRTcell.

### Data processing

The overview of the data processing used in this study is shown in Fig. S1b. Using the circular consensus sequence (CCS) reads generated from each of the sub-reads, HiFi reads, which are reads that satisfy the conditions of >3 full-pass sub-reads and QV ≥20, were automatically generated in the Sequel IIe system using SMRT Link v11.0. IsoSeq v3 (https://github.com/PacificBiosciences/IsoSeq) was used to process the HiFi reads: The 3’ and 5’ primers, concatemer and polyA tail were removed and only reads supported by at least two FLNC reads with an accuracy ≥99% were used for subsequent analysis. These reads were mapped to the human genome (hg38) using pbmm2 (https://github.com/PacificBiosciences/pbmm2) and the isoforms that were considered to be identical were combined into one according to the default value of the collapse command in IsoSeq v3.

QC assessment and filtering were performed using SQANTI3 (https://github.com/ConesaLab/SQANTI3) [59]. Human genome (hg38) and comprehensive gene annotation data from GENCODE v.41 were used as the reference for the QC. For the reference files of CAGE Peak data, polyA motif list, and polyA site data, we used the files provided in SQANTI3. Each isoform was judged against the reference to determine which of the following isoform categories it fell into: FSM, ISM, NIC, NNC, genic, intron, antisense, fusion, intergenic. Since the isoforms in each category have different probabilities of not being an artifact, a rule-based filter was applied with different conditions for each category, as shown in filtering.json (https://github.com/mihshimada/Brain_Iso-Seq). In addition, since partial RNA degradation was a concern for some of the samples in this study, additional filtering was performed for ISM isoforms in order to carefully detect ISM isoforms, i.e., only isoforms that were either determined not to be artifacts by the machine learning filter or that were detected by other long-read studies with brain samples were retained (Data provided by GTEx [20] and PacBio (https://www.pacb.com/general/data-release-alzheimer-brain-isoform-sequencing-iso-seq-dataset/)). QC was performed again on only those isoforms remaining after application of the filter.

TALON (v5.0) [60] was used for chaining multiple samples together. We initialized a TALON database using comprehensive gene annotation data from GENCODE v.41. The default settings were used as the condition for considering isoforms to be identical, except that differences in the length of the 5’ and 3’ ends of the first and last exon were allowed. We used minimap2 and samtools [61] to prepare MD-tagged SAM files for each sample and each sample was added to the data base with the conditions --cov 0.95 and --identity 0.95. Finally, only isoforms expressed in two or more samples were retained for analysis. After chaining, we focused our analysis on isoforms that were expressed in at least two samples and summarized isoforms, i.e. in this study, differences only at the both ends of the same first/last exons were not detected. We carried out all the data processing using the script ’Data_Processing1-4.py’. All command lines are described in the additional file 2.

### Rarefaction curve analysis

Rarefaction curve analysis of isoform diversity was performed to confirm whether genes or isoforms were sufficiently detected in this study. Programs subsample.py and subsample_with_category.py in cDNA_Cupcake (https://github.com/Magdoll/cDNA_Cupcake) were used for the analysis.

### PCA and cluster analysis

PCA and cluster analysis were performed using information on the presence or absence of all isoforms after filtering. The analysis was performed in R [62] and the code is described in the Additional file 3. Euclidean distance was used for PCA and clustering was performed using the Ward method.

### Gene-based comparison between brain regions

We first examined genes with high expression levels in each of the brain regions (median TPM ≥ 200) and genes expressed only in specific brain regions using the following condition: genes with TPM >2 in 3 out of 4 samples from the region of interest and TPM <1 in all other samples. Next, we examined genes with differential expression levels in different brain regions under the conditions of Wilcoxon test *P* <0.01, *t* test *P* <0.05 and fold change >5. Considering the number of samples (4 samples for the region of interest vs. 8 samples for the other regions), a threshold of *P* <0.01 was the most stringent criterion for the Wilcoxon test.

We further examined genes that have a number of isoforms (≥ 20) in each brain region and genes with a higher number of isoforms only in the region of interest compared to other regions with Wilcoxon test *P* <0.01 (4 samples for the region of interest vs. 8 samples for the other sites). Further, genes that met the following conditions were excluded: i) Genes that are not expressed except in the region of interest, ii) Genes that express only one type of isoform in the site of interest.

To explore whether isoform-rich genes were brain-specific, we performed an analysis using long-read data from GTEx (https://gtexportal.org/home/datasets). We targeted isoforms expressed in at least two of the non-brain samples, and pathway analysis was performed using genes expressing more than 20 isoforms per gene.

### Isoform-based comparison between brain regions

We examined genes with different major isoforms expressed in different brain regions. First, we determined the percentage of each isoform’s TPM relative to the total expression of the gene from which it was derived using TPM_table.py and Isoform_Proportion.py. The percentages were used to compare the cerebellum vs. the hypothalamus, the cerebellum vs. the temporal cortex, the hypothalamus vs. the temporal cortex with the condition of the median TPM of the higher expression region >20, and the median TPM of the lower expression region >5. Genes that had isoforms with Wilcoxon test *P* <0.05 and *t* test *P* <0.05 and a difference in the average TPM percentage between the two compared regions of >20% were selected. Here, a threshold of *P* <0.05 was the most stringent criterion used for the Wilcoxon test, considering the number of samples (4 samples for the region of interest vs. 4 samples for the other region). Isoforms were imaged using IGV software (v2.15.2) [63].

### Pathway analysis

Pathway analysis was performed to explore the characteristics of the genes detected in each analysis or genes to which detected isoforms belong. We used MetaCore™ software (version 6.24 build 67895, Thomson Reuters, NY, USA) with high-quality, manually curated content and the PANTHER Classification System (PANTHER 17.0) [64] to validate the results of MetaCore software. The Gene Ontology database [65, 66] was used to search for significant gene sets. We searched a maximum of the top 500 gene sets in the MetaCore analysis that met the following conditions: i) false discovery rate (FDR) <0.05 in MetaCore and replicated in PANTHER with *P* <0.05; ii) If a pathway contains fewer than 100 network objects, at least 10% of them are included in the test gene set; iii) If a pathway has more than 100 network objects, at least 10 objects are included in the test gene set; and iv) The total number of objects in the pathway is less than 3,000. When more than 10 pathways were detected, Cytoscape [67] was used to show the relationships among the top 10 pathways. The coordinates, which indicated the positional relationship between pathways, used the first and second principal components calculated based on the presence or absence of the included network objects.

### DNA methylation data

The methylation rate data obtained for the DNA derived from three brain regions of the four samples used in this study are described in detail elsewhere [43]. Briefly, DNA samples were extracted with a QIAamp DNA mini Kit (Qiagen), and DNA methylation was examined with an Infinium^®^ Methylation EPIC BeadChip kit (Illumina Inc., CA, USA). Properly filtered and normalized data were used for the analysis. Reference-free estimation of the percentage of cell components was performed using MeDeCom [38].

### Integrative analysis of transcriptome and methylation data

We examined whether there were any methylation sites with different methylation rates between brain regions in the vicinity of genes that expressed different major isoforms among regions. We first detected methylation sites with different methylation rates between brain regions (differentially methylated position, DMP). For each comparison of the cerebellum vs. the hypothalamus, the cerebellum vs. the temporal cortex, and the hypothalamus vs. the temporal cortex, methylation sites with Wilcoxon’s test *P* <0.05 and *t* test *P* <0.05, and with >20% difference in methylation rate between groups were identified. We then compared the presence of the DMPs in the vicinity of genes, between genes expressing different major isoforms, and genes expressed in all brains using CpG_annotation.py. Annotation information such as whether the methylation site is upstream or downstream of the gene was obtained from the annotation file provided by Illumina Inc. (infinium-methylationepic-v-1-0-b5-manifest-file.csv).

Further, we determined whether the differences between isoforms corresponded to different first and/or last exon, exon with different 5′ and/or 3′ splice site, intron retention, exon skipping, cassette exon, or others. First, isoform pairs that were differentially expressed between regions were identified. That is, if an isoform was detected as being differentially expressed between the regions, meeting the conditions described above, and it was the most major isoform at one of the regions being compared, then the comparison was made with the most major isoform at the other site (Fig.5b, A). If the most major isoform was common to both regions and there was a difference in expression of the second most major isoform at one region, then the second most major isoform at the other region was also compared (Fig.5b, B). Similarly, in other cases, lower ranking isoforms were compared to each other (Fig.5b, C). Next, we identified areas in the vicinity of isoforms that differed from each other according to their structural patterns (Fig. S13). Detailed definitions of the areas are shown in Fig. S13. Comparison of isoforms and extraction of relevant areas was performed with AS_investigation.py. We then searched for DMPs of brain regions in these areas. As a control for comparison, we performed the same analysis between isoforms that showed no differences in expression patterns between brain regions to search for the extent to which DMPs were present in areas associated with differences in isoforms. Genes with TPM >10 were targeted, and of the isoforms for which *P* >0.2 in any of the statistical tests of comparisons between brain regions, isoforms that were ranked second or higher in terms of their expression level within a gene at any region were included. Comparisons of these selected isoforms within the same gene were used as controls for statistical analysis.

### Utilization of proteome analysis for the validation of novel isoforms

To confirm whether the newly identified isoform is translated and exists as a protein in the brain, we utilized the results of LC-MS/MS analysis of human postmortem cortex obtained in a previous study [40]. The raw data was downloaded from the ProteomeXchange Consortium (https://www.proteomexchange.org) and processed as an 11-plex tandem mass tag using MaxQuant [68]. TMT parameter files (Lot No SI258088 and SJ258847) were obtained from Thermo Fisher Scientific website. The ’Type’ for the group-specific parameters was set to ‘11plex TMT’, and the charge for each was set to the value in the TMT parameter files. In addition to Trypsin, LysC was also selected for ‘Digestion’. For the ’Sequence’ in the global parameters, we used the GENCODE protein-coding transcript translation sequences Release41, which is the same version of the transcript information used for identifying isoforms. All other settings were left as default values. Only the peptides used for identifying proteins detected with a criterion of 1% FDR were analyzed. We also conducted an analysis with the same settings but changed the amino acid sequences of the predicted ORF for the novel isoforms to the reference only. We then searched for peptides that did not match the amino acid sequences registered in GENCODE and those estimated from the known transcripts identified in our study, specifically targeting those that were detected for the amino acid sequences derived from the novel isoforms.

### Bioinformatics workflow and command lines

The command lines used in this study are listed in Additional file 2 and 3 (R code). The python programs and json file are also available on GitHub (https://github.com/mihshimada/Brain_Iso-Seq)

## List of abbreviations

Ce: cerebellum
Hy: hypothalamus
Tc: temporal cortex
FSM: full splice match
ISM: incomplete splice match
NIC: novel in catalog
NNC: novel not in catalog
DMP: differentially methylated position

## Declarations

### Ethics approval and consent to participate

Written informed consent was obtained from all participants or their families, and the protocols were approved by the ethics committees of all the collaborative institutes.

### Consent for publication

Not applicable.

### Availability of data and materials

Iso-Seq analysis data has been deposited into DDBJ (https://www.ddbj.nig.ac.jp/) with the BioProject accession number: PRJDB15555.

### Authors’ contributions

MS designed the study. AK and RG collected and dissected brain samples. MS and YO carried out experiments and data analyses. TM, MH, AF and KT interpreted the results. MS drafted the manuscript, and all authors contributed to the final version of the paper.

## Acknowledgements

We extend our gratitude to all the donors who provided samples for this study. The Genotype-Tissue Expression (GTEx) Project was supported by the Common Fund of the Office of the Director of the National Institutes of Health, and by NCI, NHGRI, NHLBI, NIDA, NIMH, and NINDS. The data used for the analyses described in this manuscript were obtained from the GTEx Portal on 08/15/2022.

