## Additional_file1 for "Identification of Region-Specific Gene Isoforms in the Human Brain Using Long-Read Transcriptome Sequencing and Their Correlation with DNA Methylation"

Fig. S1. Graphical abstract and analytical flow of Iso-seq data

Fig. S2. Quality check results for Iso-seq raw data

Fig. S3. Indicators of good quality in the Iso-seq analysis

Fig. S4. Rarefaction curves for each isoform category by brain region

Fig. S5. Relationships between samples inferred from Iso-Seq and genome-wide methylation data

Fig. S6. Examination of expression (TPM) features

Fig. S7. Results for isoform length

Fig. S8. Frequency distribution of exons in each isoform category

Fig. S9. The examples of the novel isoforms validated by LC-MS/MS

Fig. S10. Pathway analysis of genes associated with gene expression levels

Fig. S11. Pathway analysis of genes associated with genes with many isoforms

Fig. S12. Relationship between expression level and the number of isoforms per gene

Fig. S13. Definition of isoform differences and related areas

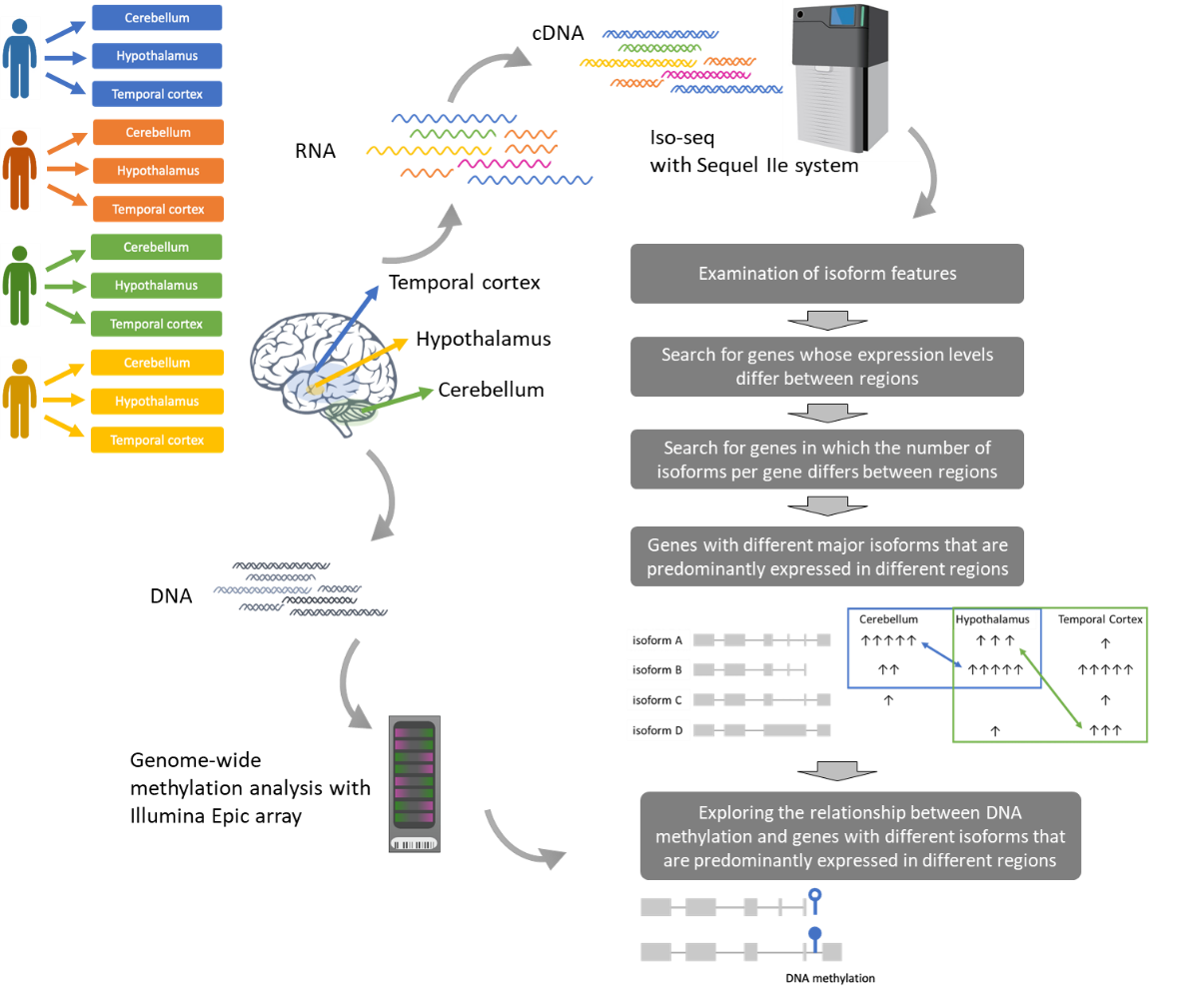
**Fig. S1.**
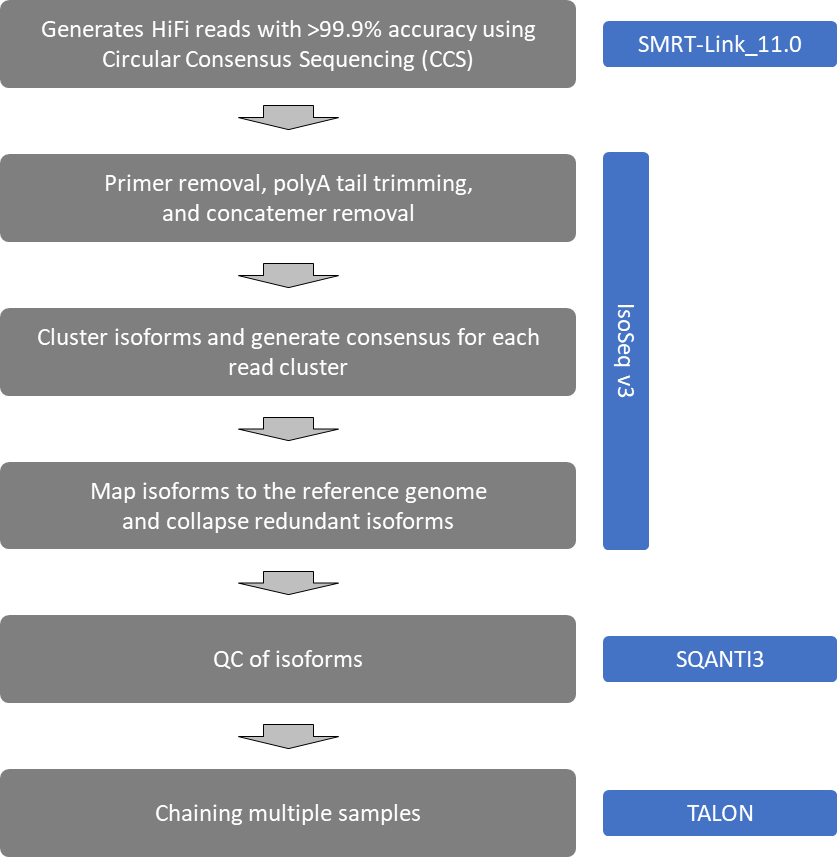
**Graphical abstract and analytical flow of Iso-seq data. a** Tissues were excised from three brain regions, cerebellum (Ce), hypothalamus (Hy), and temporal cortex (Tc) from the same samples, RNA was extracted and Iso-seq was performed. DNA methylation was evaluated using DNA from the same tissues. **b** Analysis flow of Iso-seq data and software used.

**b**

**a**

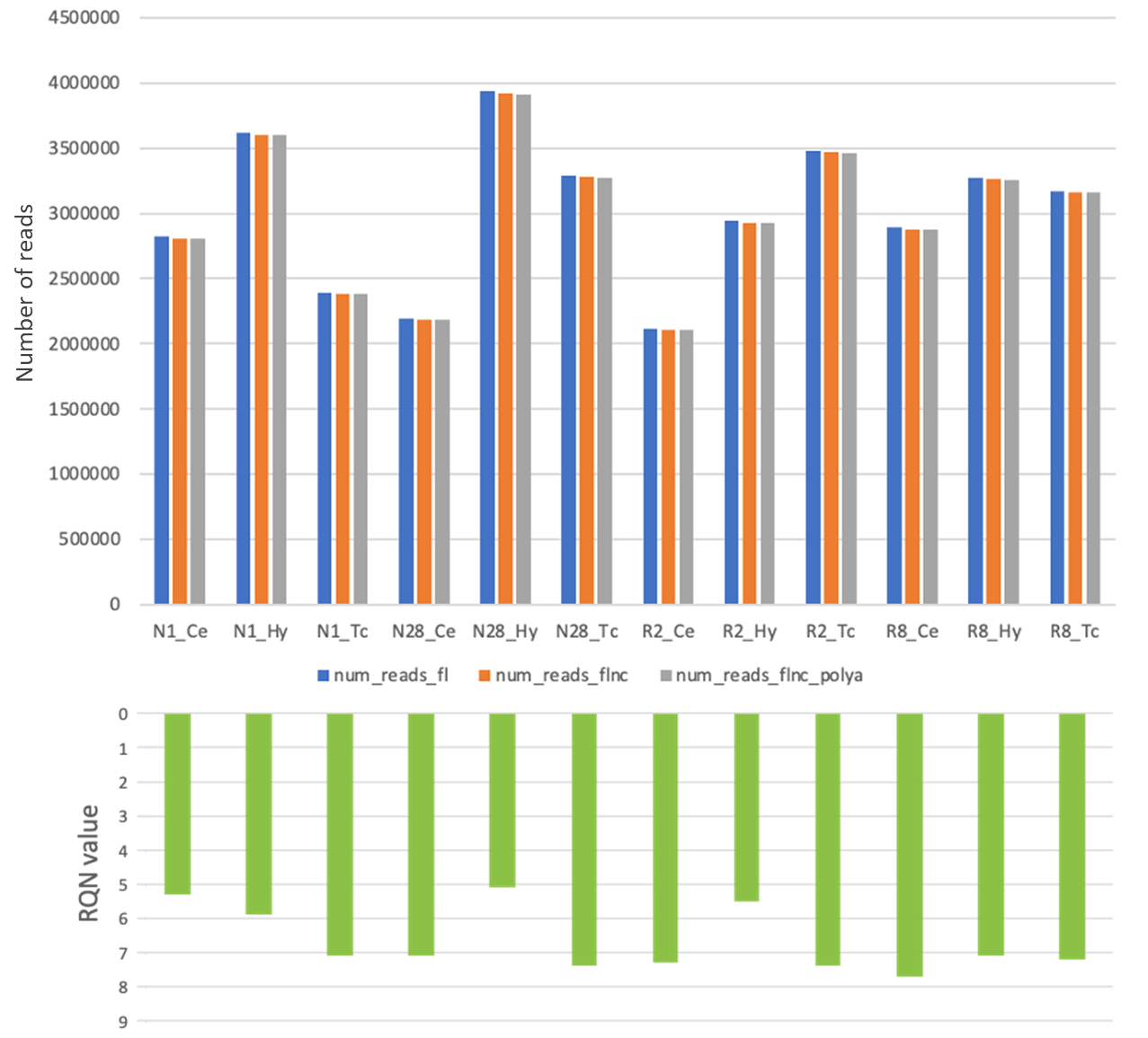

**Fig. S2. Quality check results for Iso-seq raw data.** Numbers of full-length (fl) reads, full-length non-concatemer (flnc) reads, and flnc reads with polyA tails. The relationship between the number of each read and the RQN value for RNA quality is also shown; no relationship is observed between the RQN and the number of detected reads.

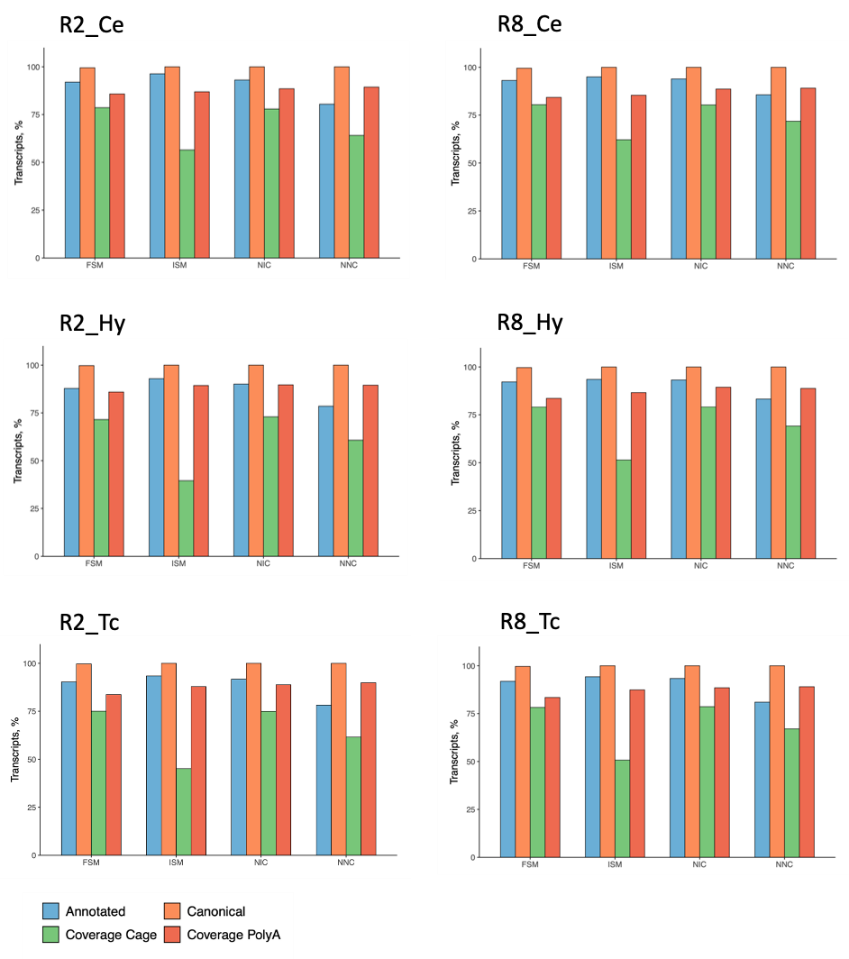

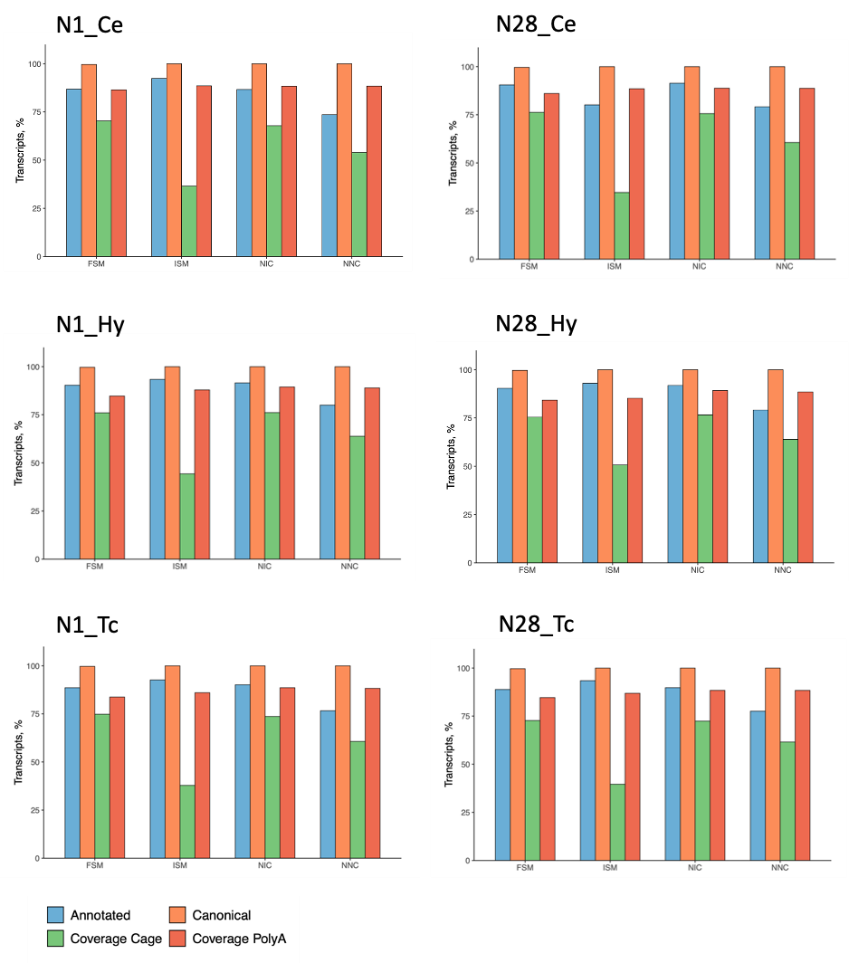

**Fig. S3. Indicators of good quality in the Iso-Seq analysis.**  “Annotated” indicates that the transcription start site (TSS) of each isoform is less than 50 bp from the TSS of a known gene. “Canonical” indicates isoforms with canonical junctions using the following splice donor and acceptor sites: GT/AT, GT/AG, GC/AG, and AT/AC. “Coverage Cage” indicates whether there is a cage peak within 10 kb of the TSS. “Coverage PolyA” indicates whether there is a polyA tail. Although the "Coverage Cage" of ISM is low, the remainder of the indicators are generally high.

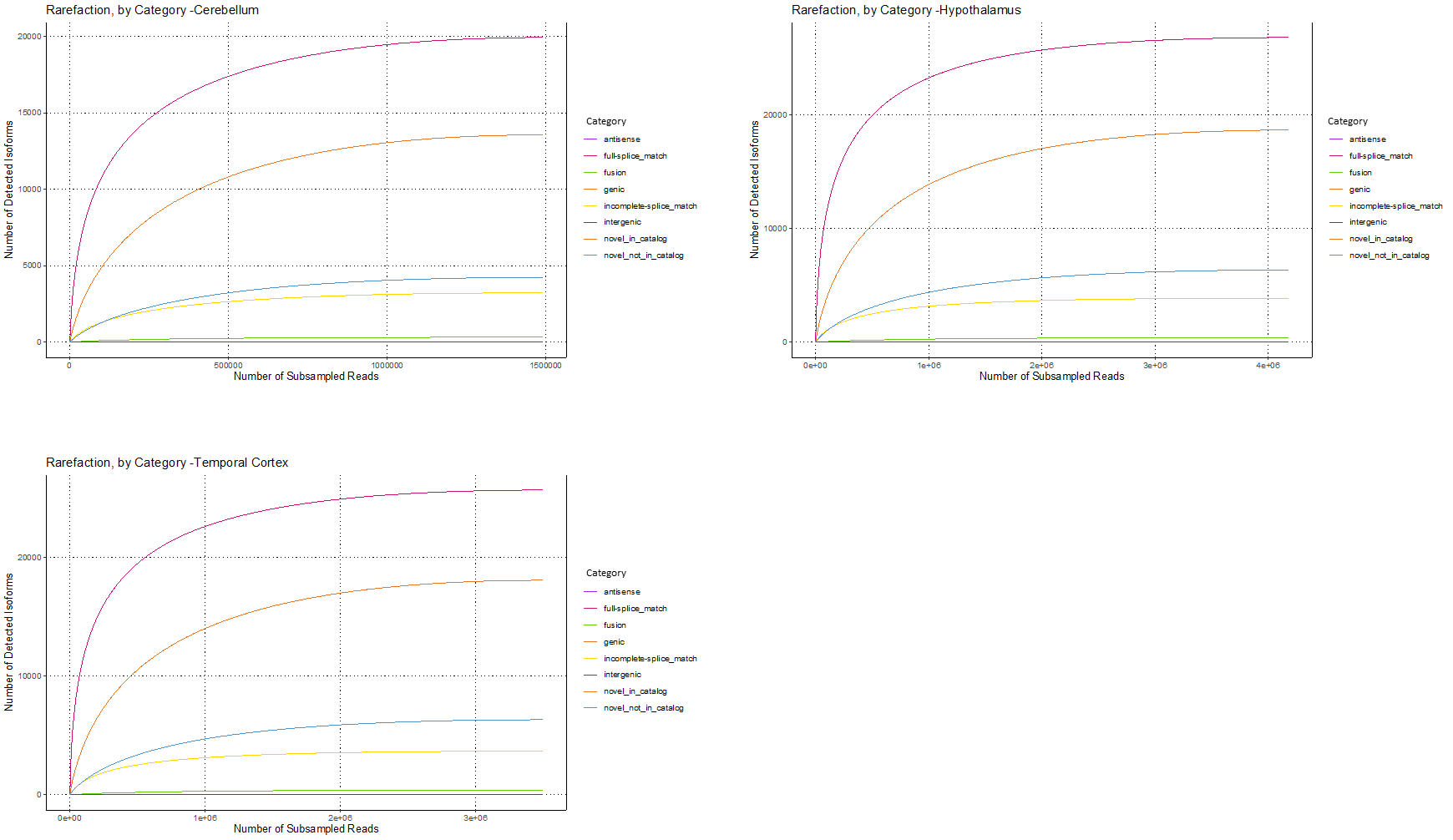
**Fig. S4. Rarefaction curves for each isoform category by brain region. a** Cerebellum. **b** Hypothalamus. **c** Temporal cortex

**c**

**b**

**a**

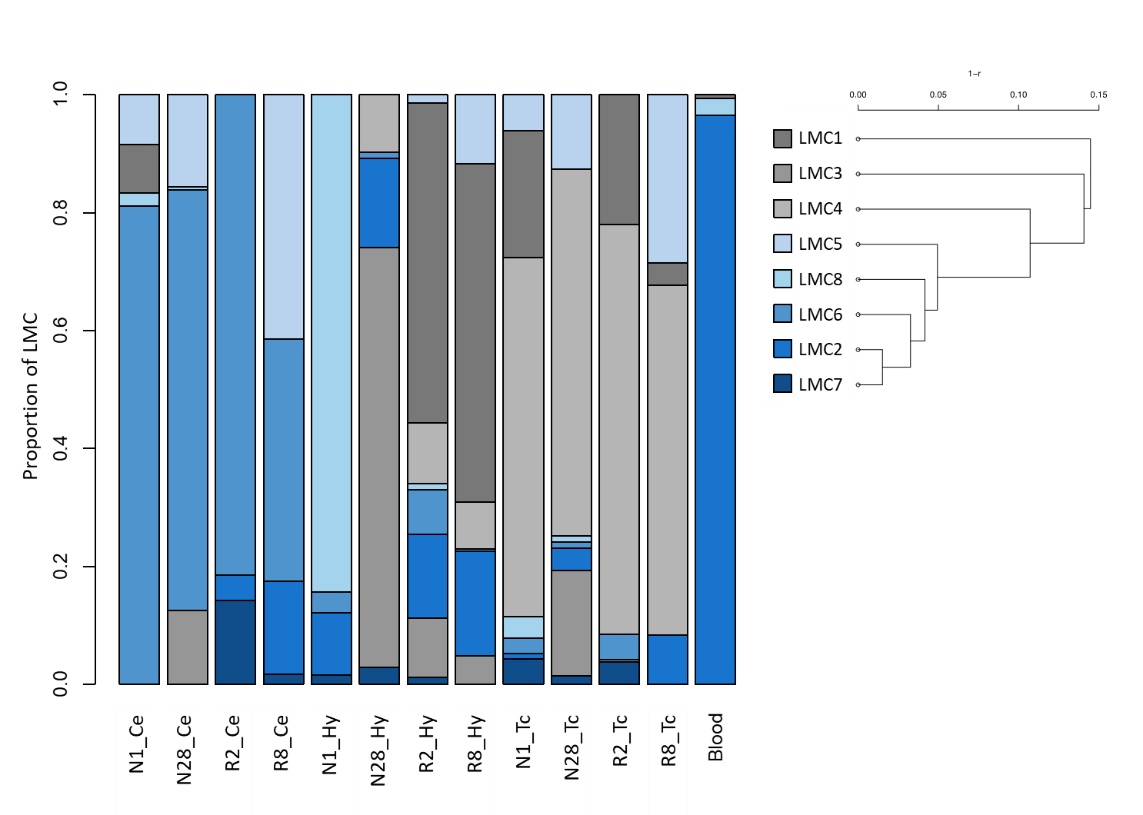

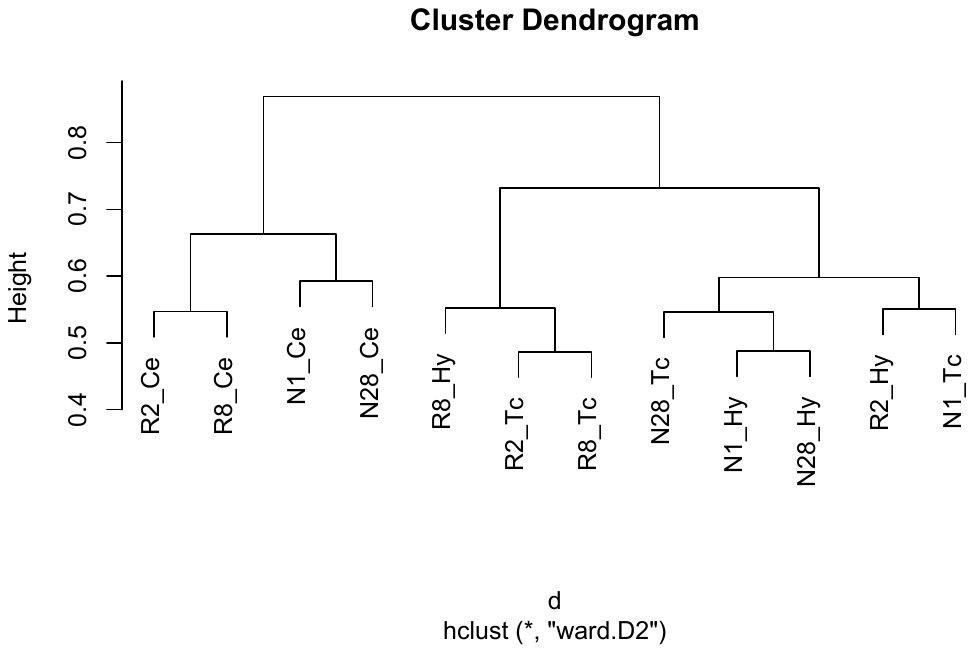

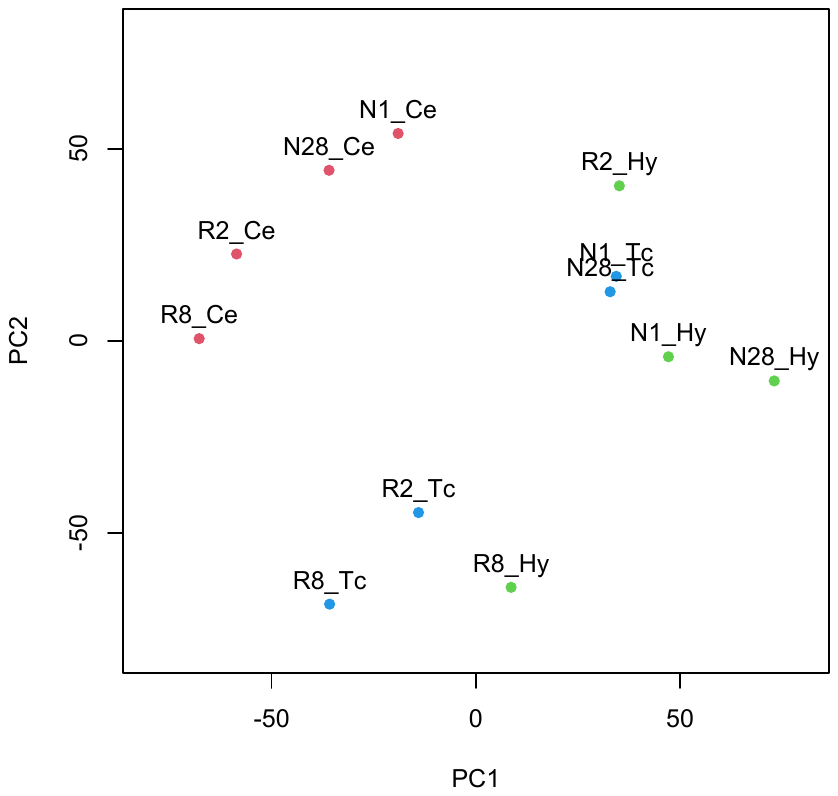
**Fig. S5. Relationships between samples inferred from Iso-seq and genome-wide methylation data.** **a** Principal component analysis based on isoforms identified by Iso-Seq. **b** Cluster analysis based on Iso-Seq results. **c** Reference-free estimates of cell composition ratios using genome-wide methylation data. Latent methylation component (LMC) corresponds to the cell type assumed to be included in the analysis. One example of peripheral blood-derived DNA methylation data is included as a control, and the ratio of LMC in each sample is shown (by color) in order of proximity of the branches according to the cluster analysis on each LMC.

**c**

**b**

**a**

**a**

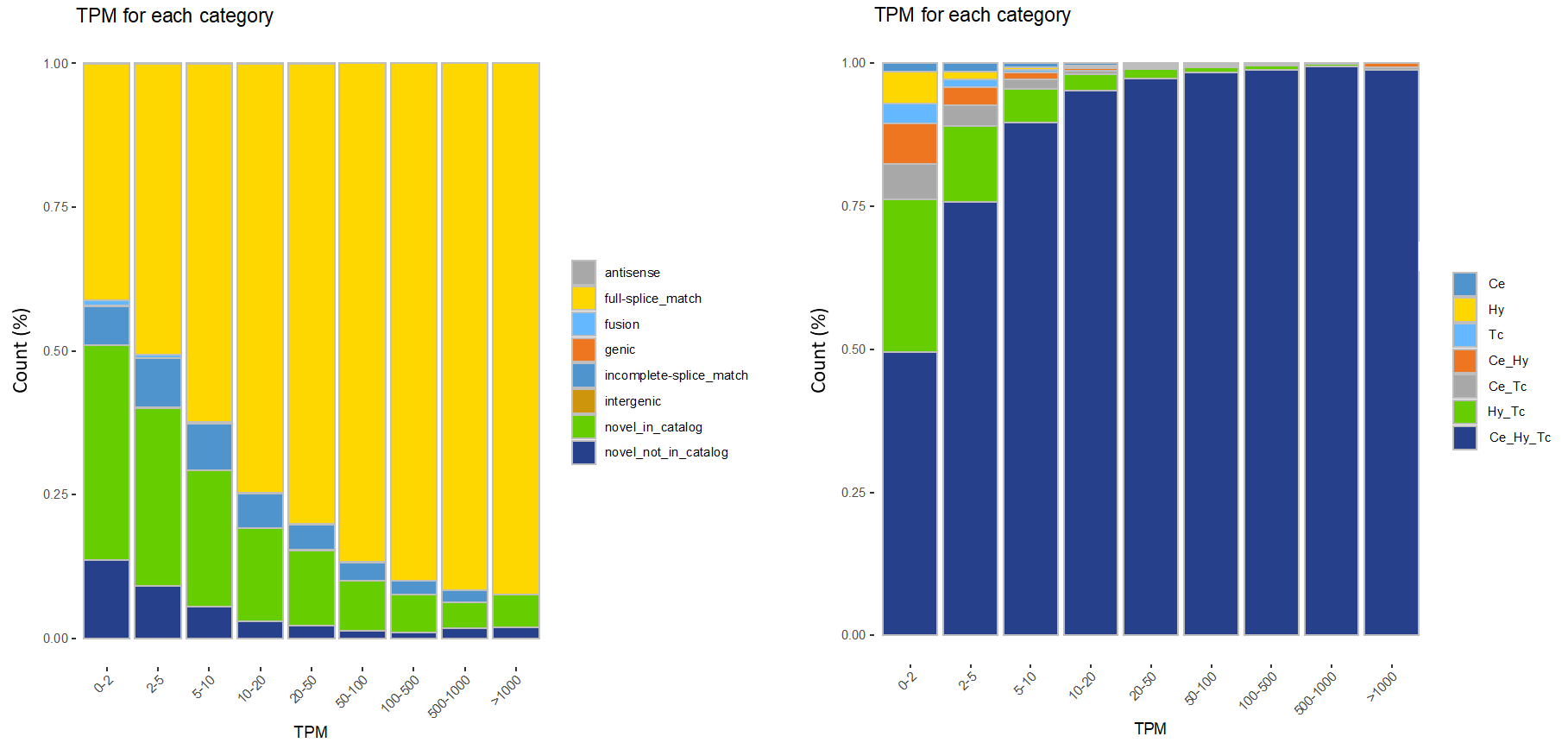
**Fig. S6. Examination of expression (TPM) features. a** Percentage of isoform categories in each TPM category. While the proportion of FSM increases as TPM increases, the proportion of novel isoforms decreases. **b** Proportion of isoforms expressed in different brain regions in each TPM category. While the proportion of isoforms expressed in all the three regions increases as TPM increases, the proportion of isoforms expressed in up to two regions decreases.

**b**

**a**

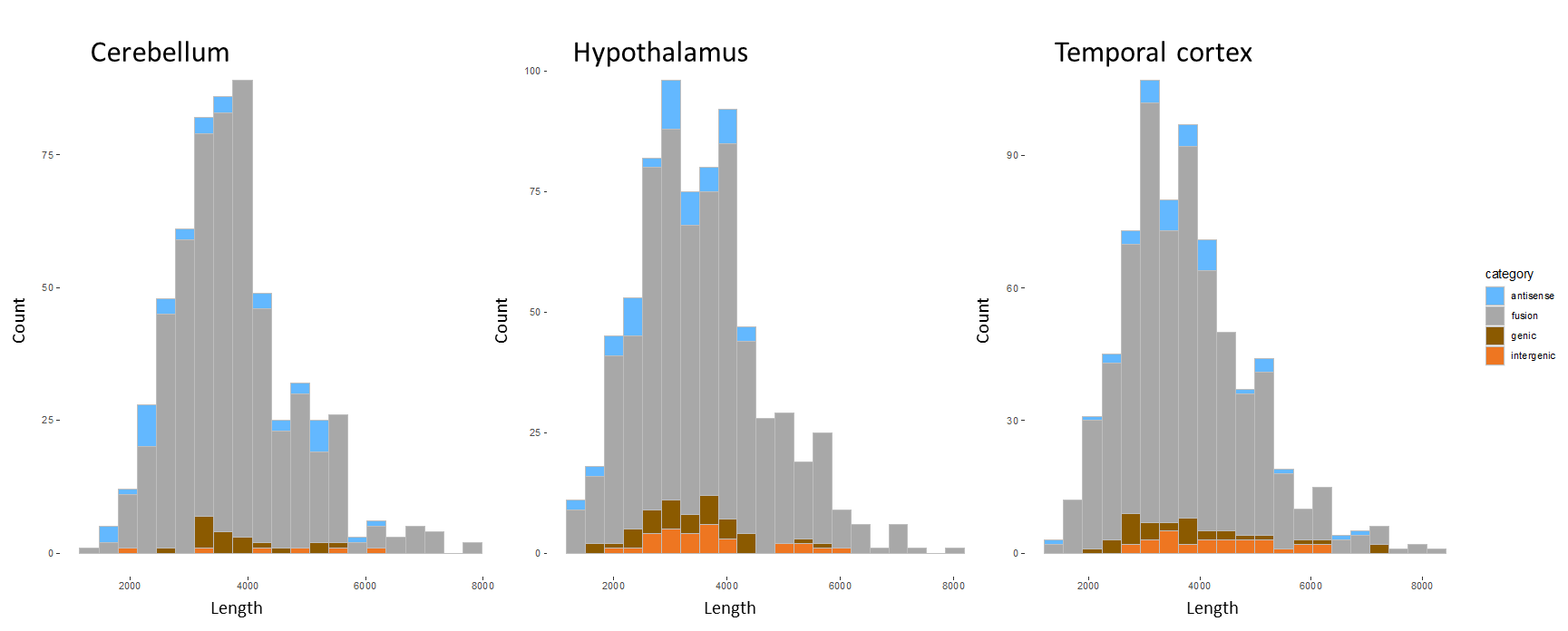

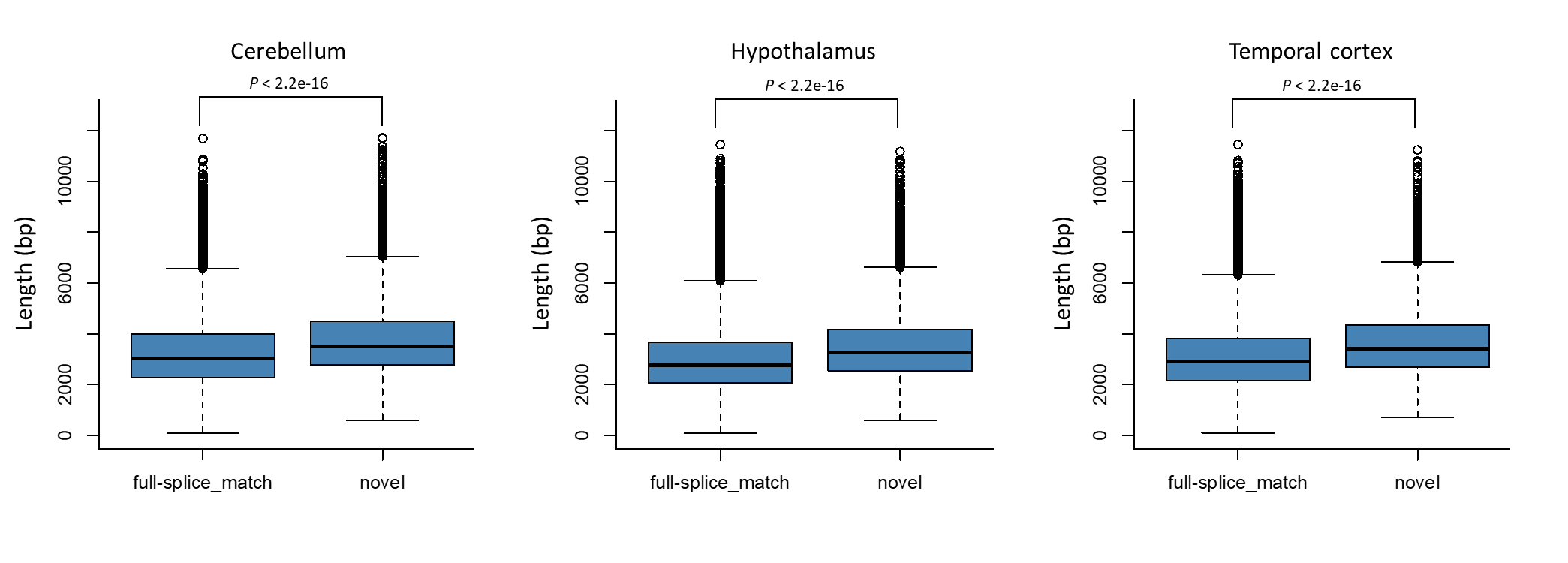

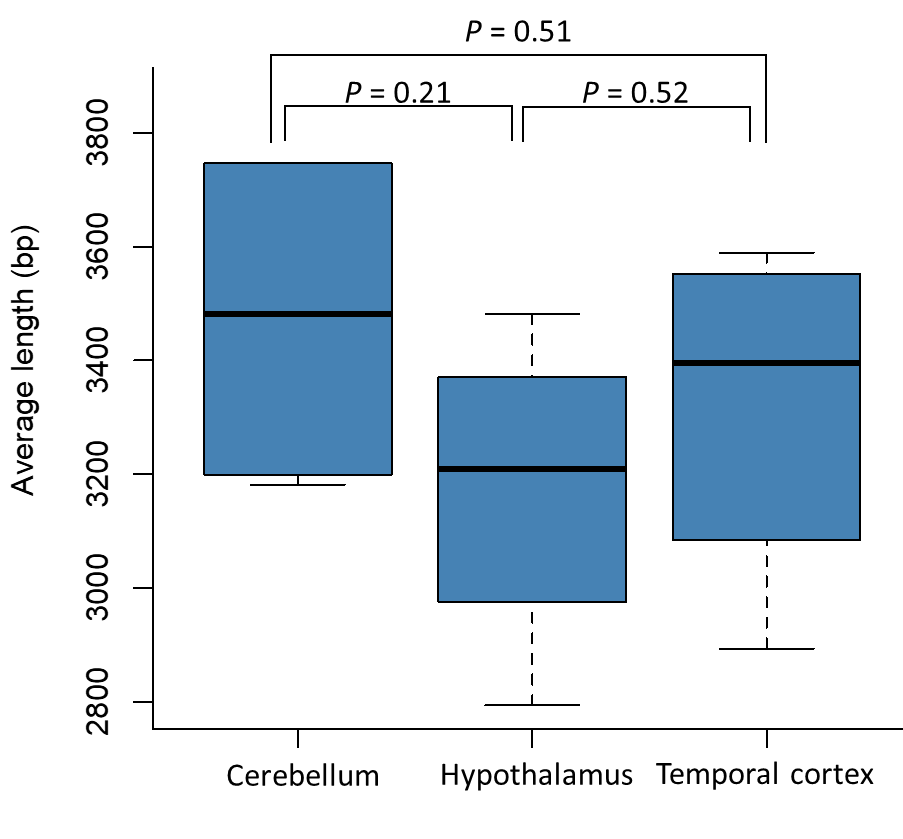

**c**

**b**

**a**

**Fig. S7. Results for isoform length. a** Comparison of lengths per sample between the brain regions. **b** Length distribution of isoform categories with few detections. **c** Comparison of FSM and new isoform lengths by region.

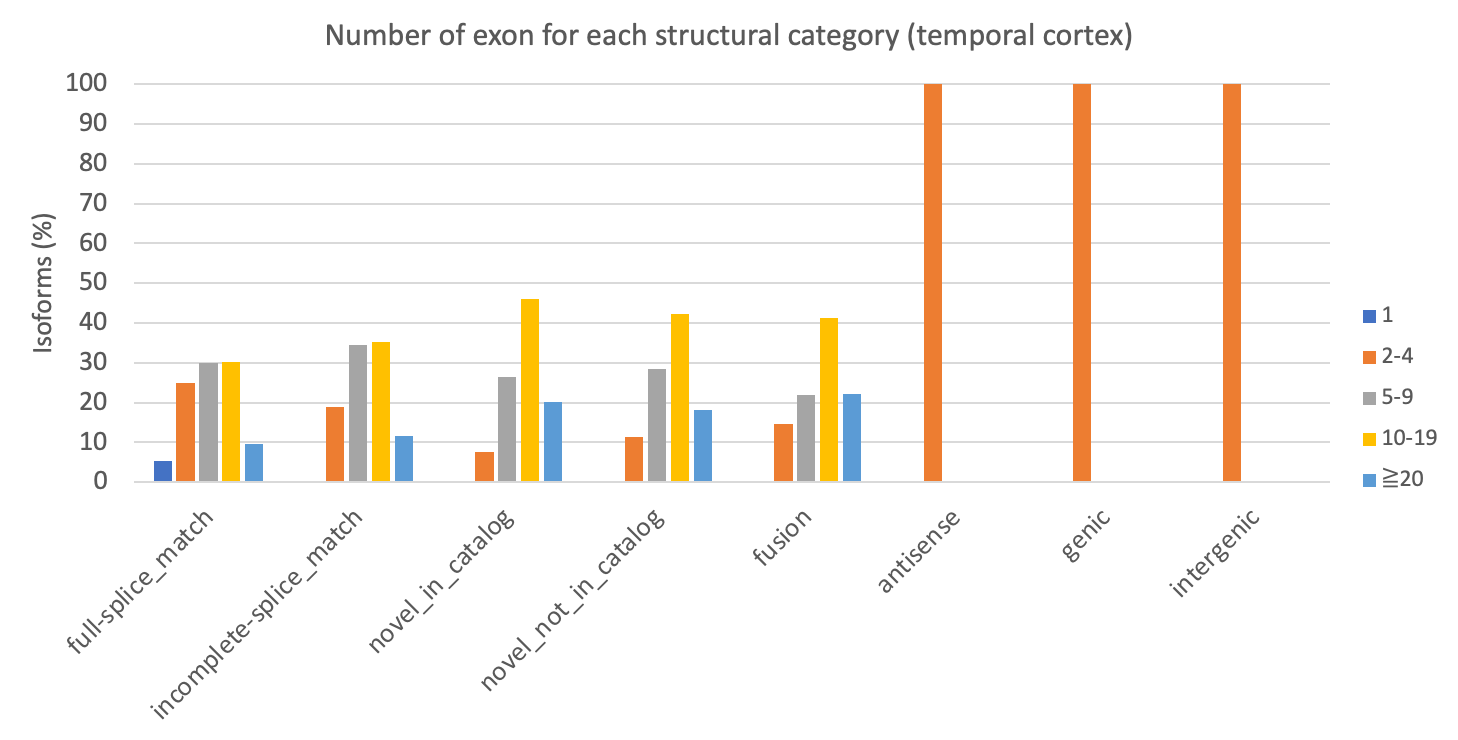

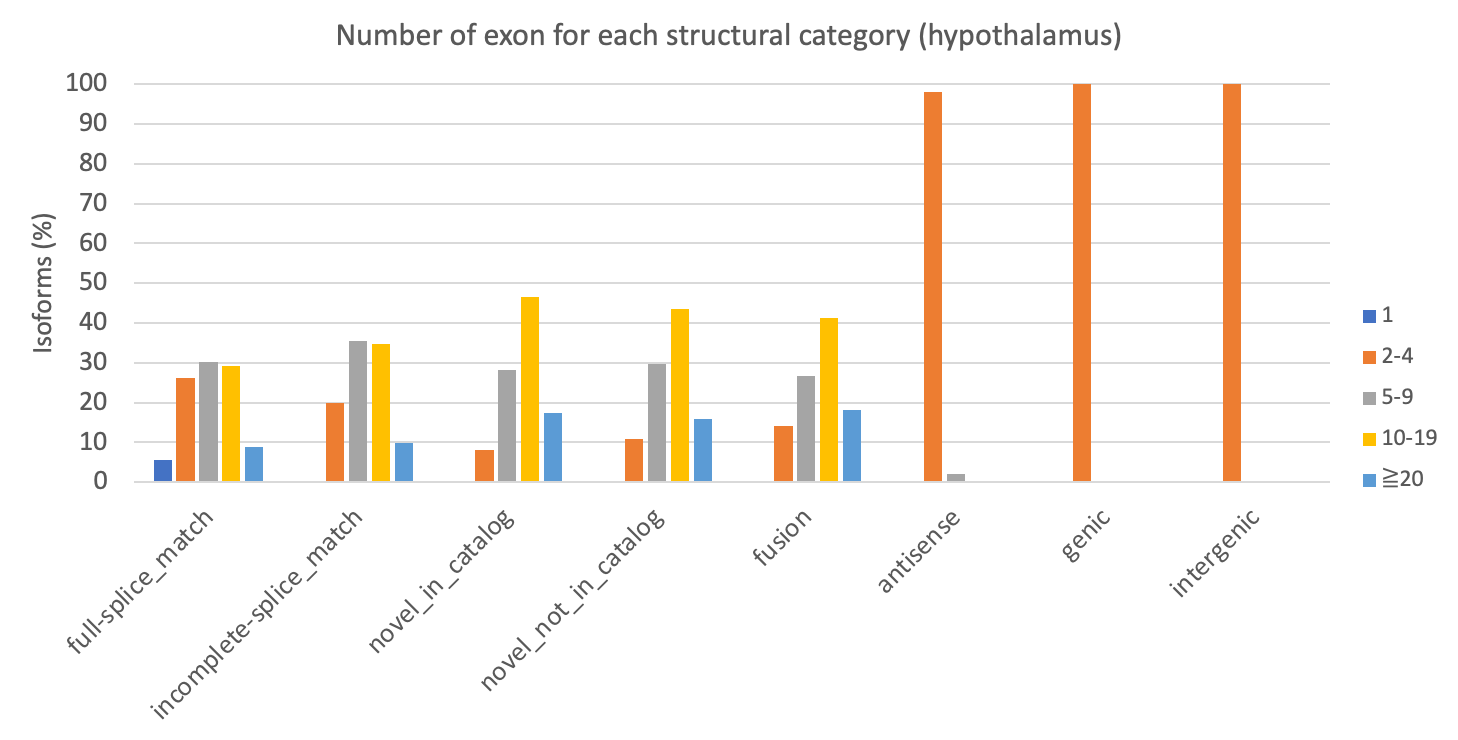

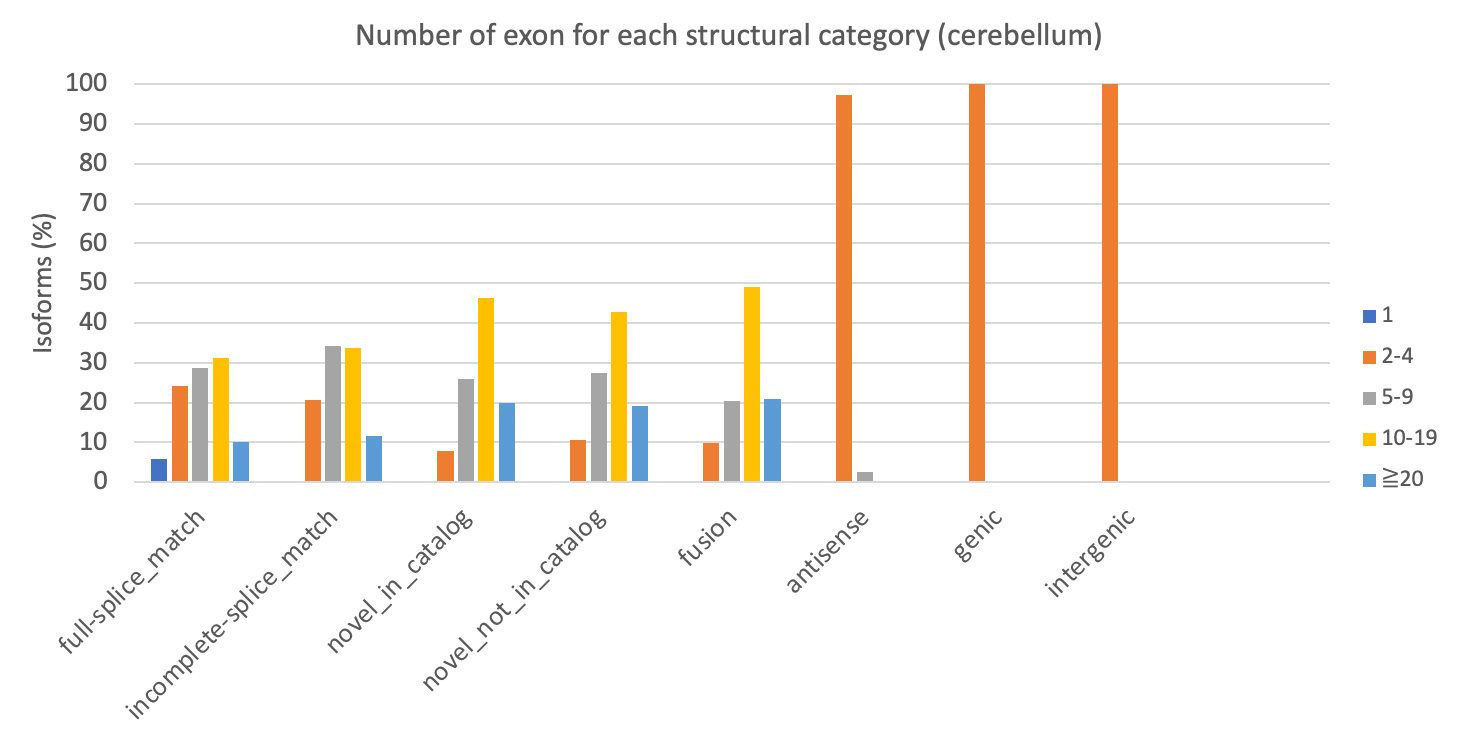
 **Fig. S8. Frequency distribution of exons in each isoform category. a** Cerebellum, **b** Hypothalamus, and **c** Temporal cortex. In all regions, NIC and NNC had more exons than FSM and ISM, with more isoforms in the 10–19 category; antisense, genic, and intergenic isoforms had fewer exons and were mostly classified in the 2–4 category.

**c**
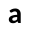

**b**
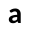

**a**

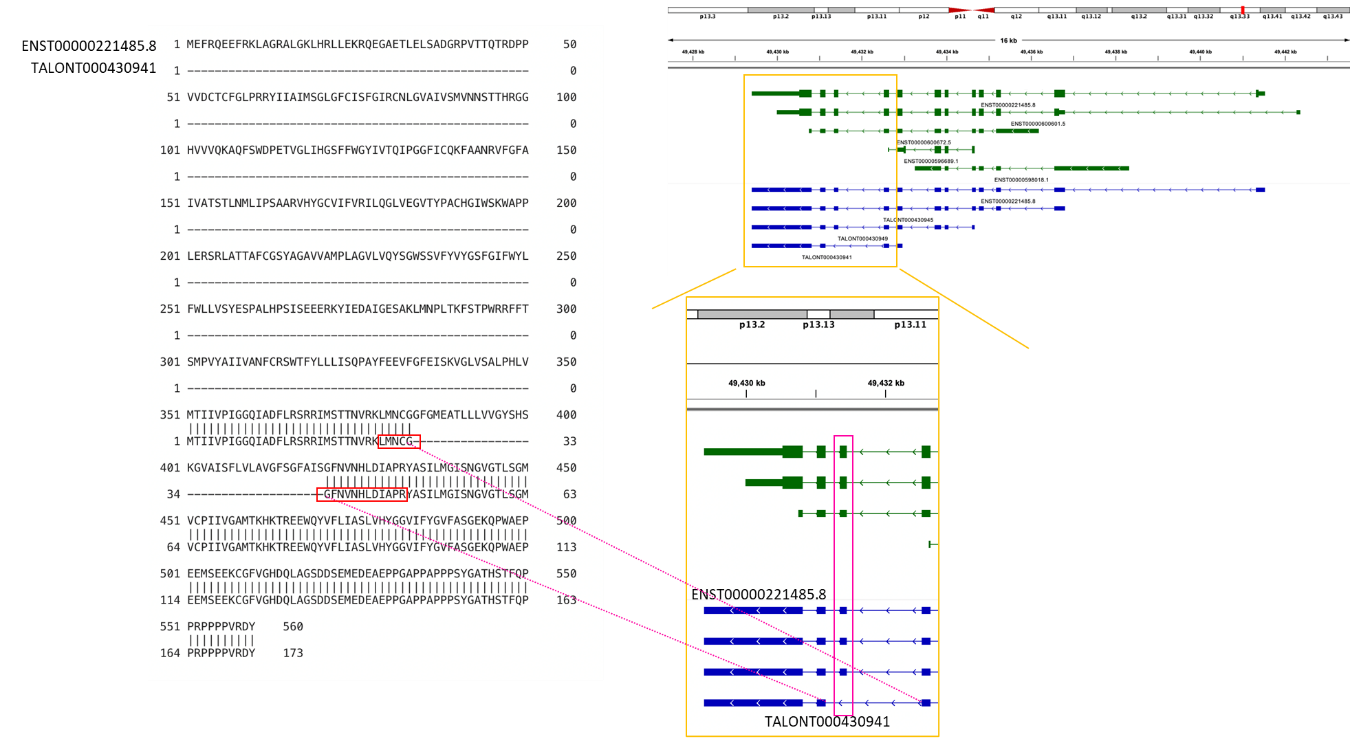

**b**

**a**

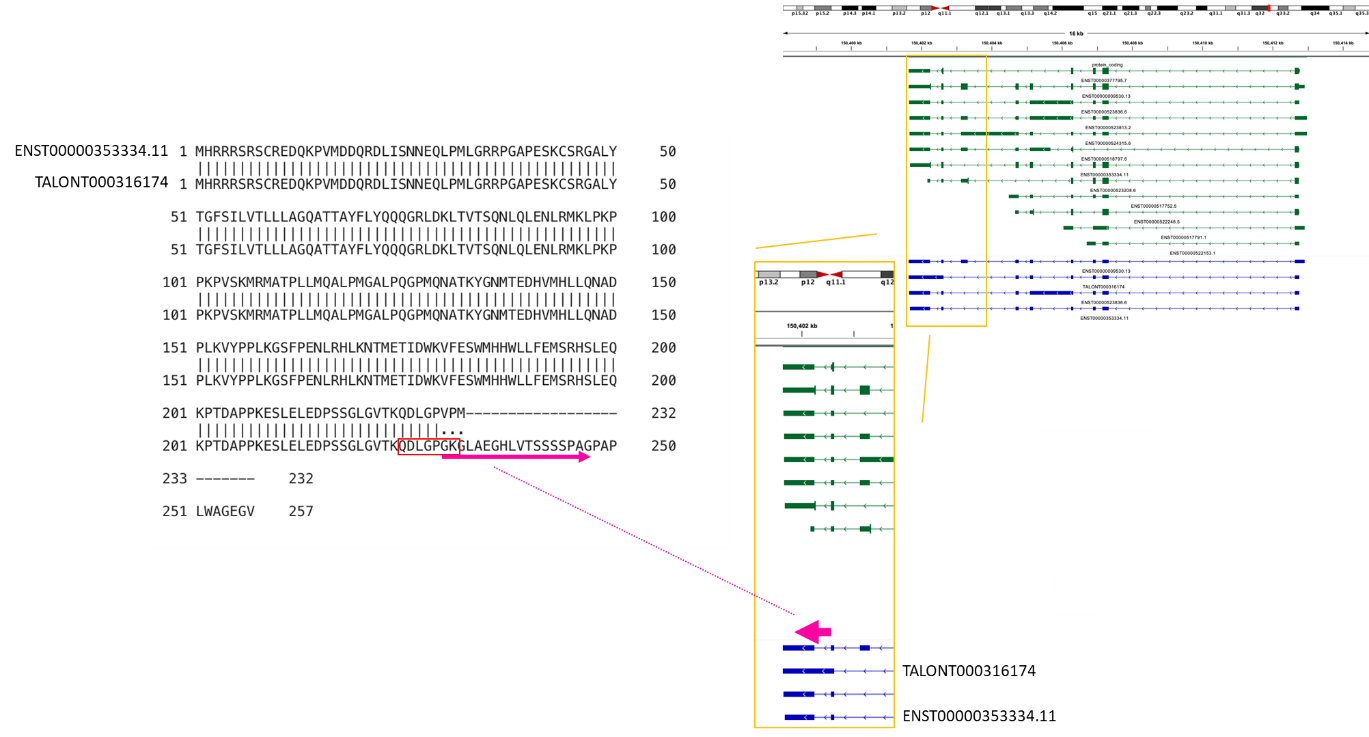

**c**

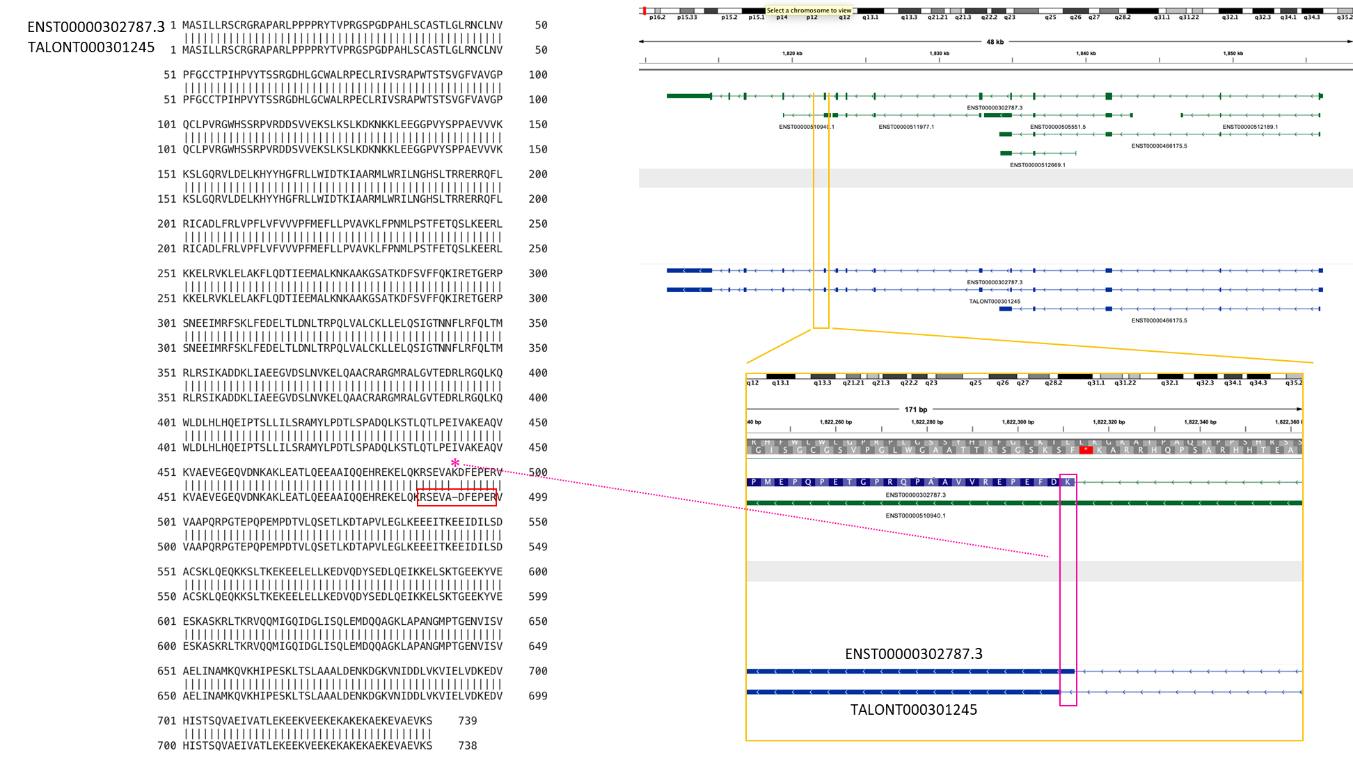

**Fig. S9. The examples of the novel isoforms validated by LC-MS/MS.** Each blue isoform represents the isoform detected in this Iso-Seq analysis, while the green isoforms represent all isoforms registered in GENCODE v.41. The amino acid sequence alignment was performed using EMBOSS Needle (https://www.ebi.ac.uk/Tools/psa/emboss_needle/). **a** An example of exon skipping. In the novel isoform, TALONT000430941, of the *SLC17A7* (Solute Carrier Family 17 Member 7) region, the third exon from the last exon was skipped, and the amino acid sequence resulting from the joining of the exons before and after that exon was detected. **b** An example of intron retention. In the novel isoform, TALONT000316174, of the *CD74* (CD74 Molecule, Major Histocompatibility Complex, Class II Invariant Chain) region, the intron between the last exon and the exon before it was retained, and the translated amino acid sequence resulting from the retained intronic region was detected. **c** An example of novel splice junction of NNC. In the novel amino acid sequence (TALONT000301245) of the *LETM1* (Leucine Zipper And EF-Hand Containing Transmembrane Protein 1) region, a one-amino-acid shift was found in the 5' splice site of the 10th exon. The detection of the amino acid sequence containing this amino acid indicated the presence of a novel splice junction.

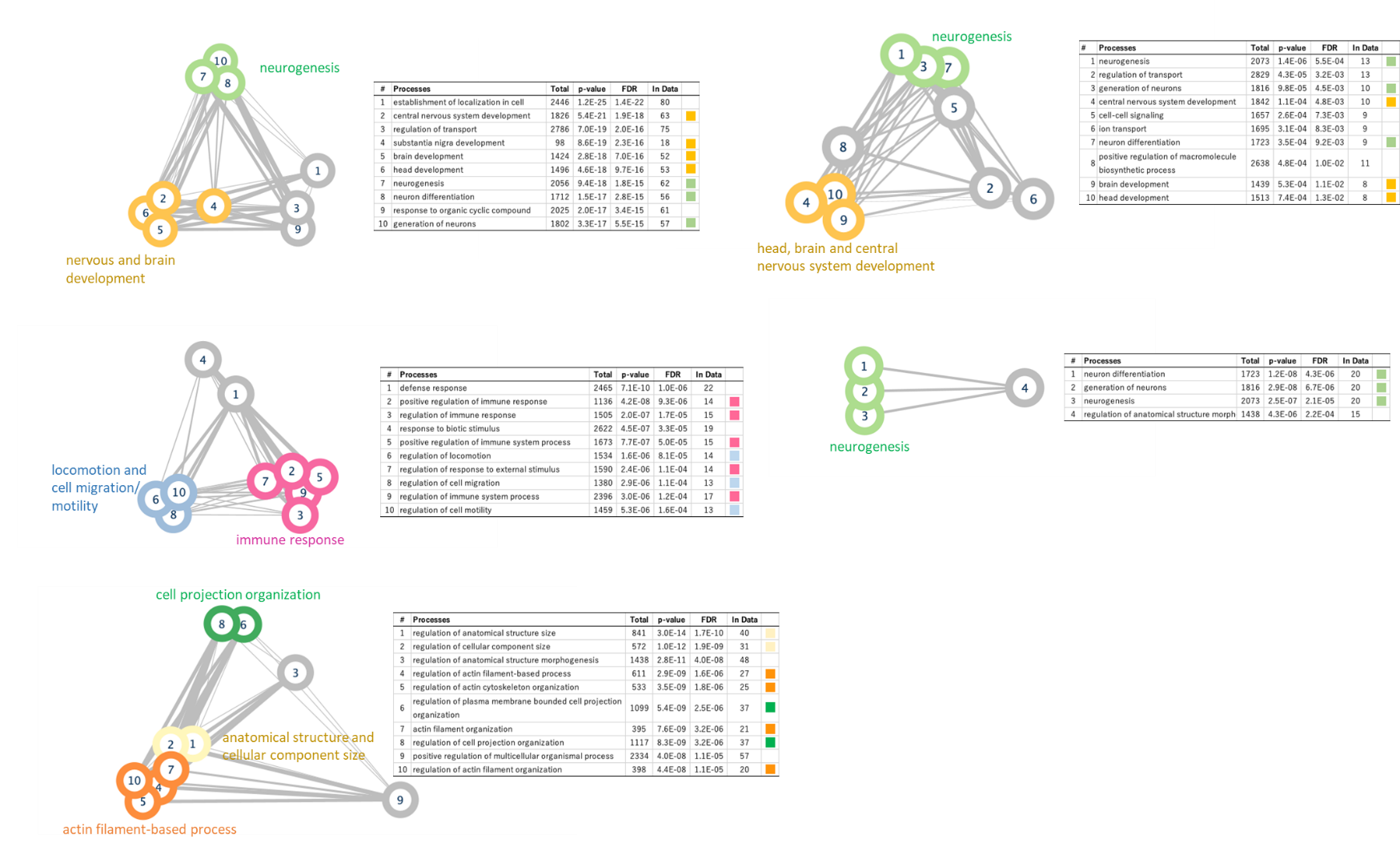
**Fig. S10. Pathway analysis of genes associated with gene expression levels.** When more than 10 pathways are detected, the top 10 pathways with the smallest FDR are shown. Edge thickness is proportional to the number of related genes shared among pathways. **a** Pathway analysis results for genes with high expression (transcripts per million (TPM) >200) in all brain regions. **b** Pathway analysis results for genes that are upregulated in the cerebellum. **c** Pathway analysis results for genes that are upregulated in the hypothalamus. **d** Pathway analysis results for genes that are upregulated in the temporal cortex. **e** Pathway analysis results for genes that are downregulated in the cerebellum.

**d**

**c**

**e**

**b**

**a**

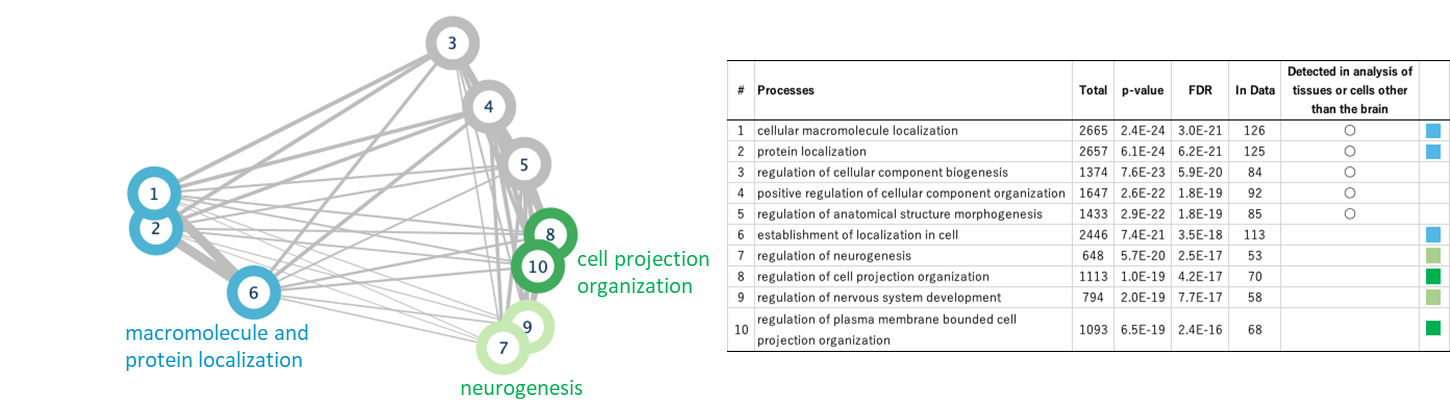

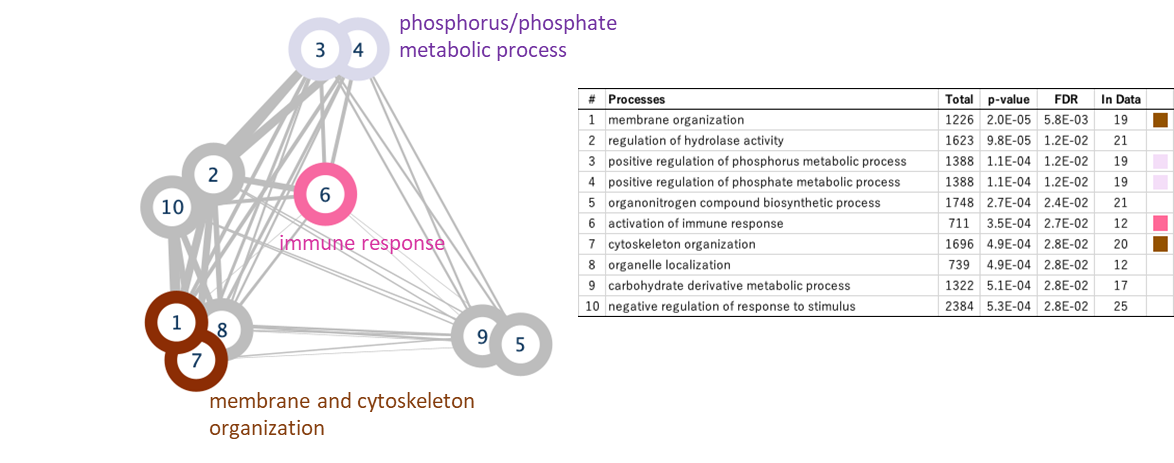

**b**

**a**

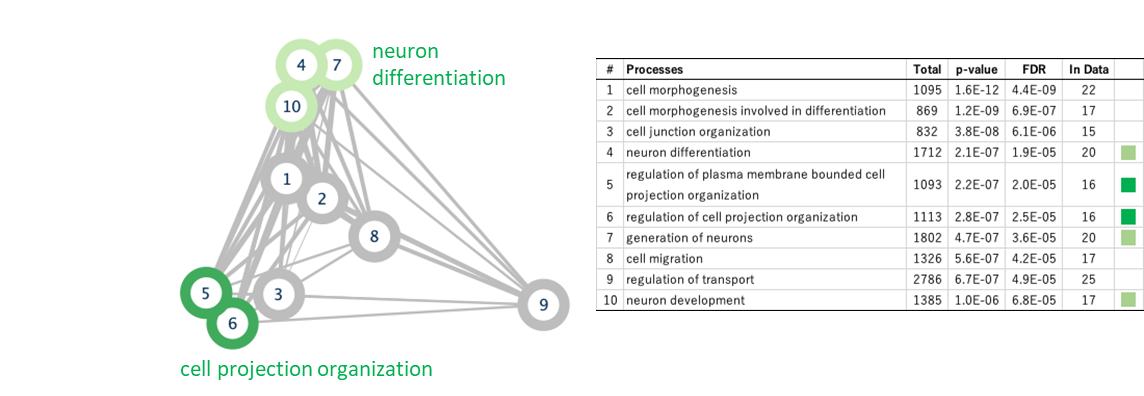

**c**

**Fig. S11. Pathway analysis of isoform-rich genes.** When more than 10 pathways are detected, the top 10 pathways with the smallest FDR are shown. Edge thickness is proportional to the number of related genes shared among pathways. No significant pathways were detected in the pathway analysis using isoform-rich genes in the cerebellum. **a** Results of isoform-rich genes in all of the brain regions. **b** Results of isoform-rich genes in the hypothalamus. **c** Results of isoform-rich genes in the temporal cortex.

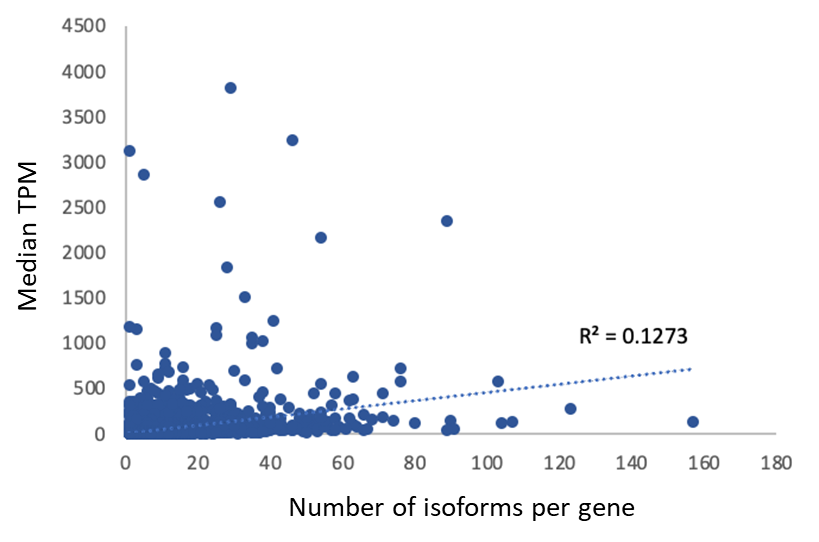

**Fig. S12. Relationship between expression level and the number of isoforms per gene.** The expression levels (TPM) and the number of isoforms per gene are weakly correlated with a correlation coefficient of 0.127.

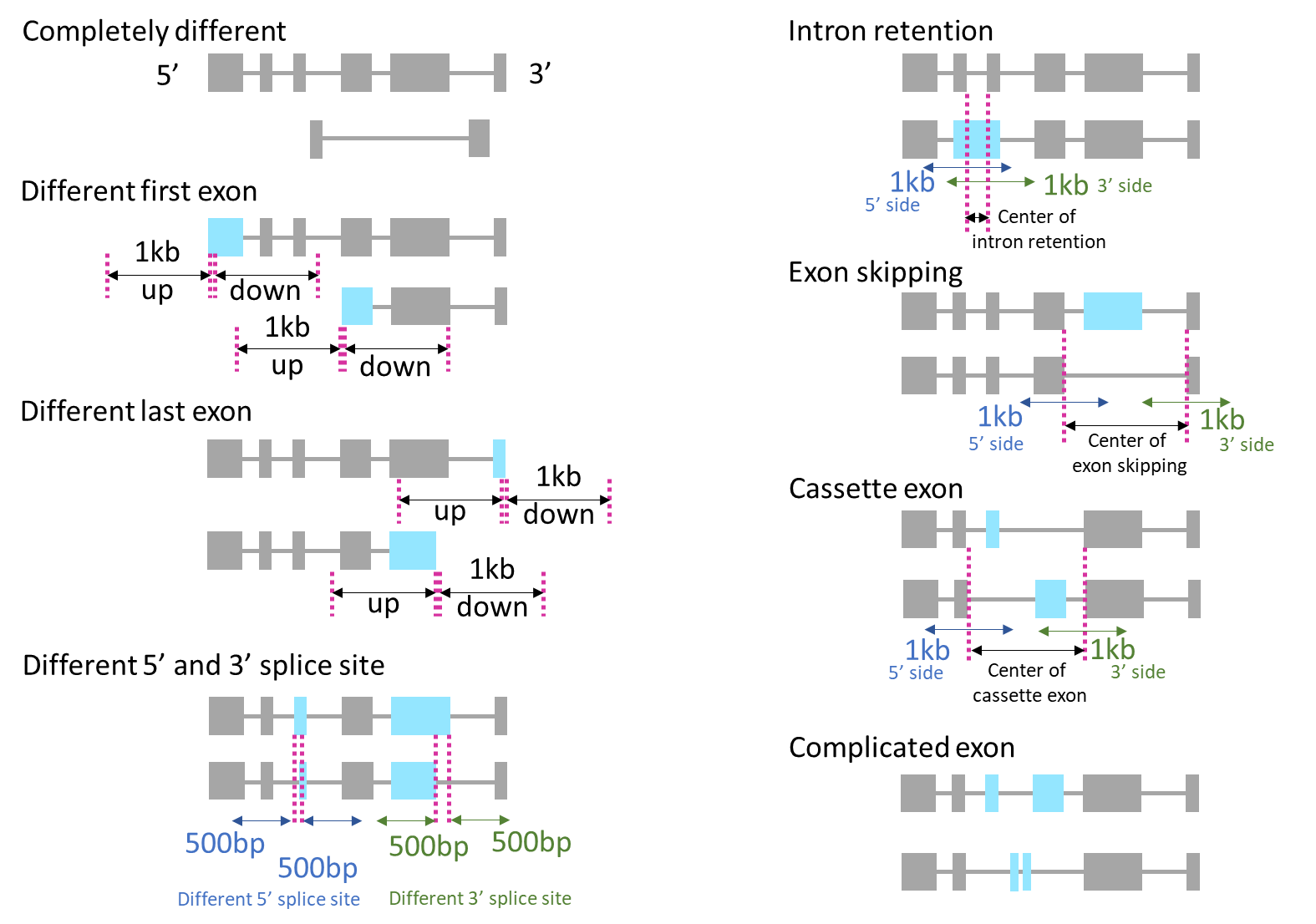
**Fig. S13. Definitions of isoform differences and related areas.** We examined whether or not there is a difference between each pair of the compared isoforms. “**Completely different**” is a case in which exons do not overlap at all. Here, if the shorter of the two exons to be compared overlapped by more than 50%, they were considered to be the same exon. “**Different first exon**” and “**Different last exon**” indicate that the first exon or the last exon are different, and the region 1 kb upstream and 1 kb downstream from either the TSSs or the transcription termination sites were defined as relevant regions, respectively. “**Different 5′ and 3′ splice site**” are cases where the 3' and/or 5' sides of overlapping exons do not match. The 5' end of the first exon and the 3' end of the last exon were excluded from the analysis. For “**Intron retention**”, the region of the expressed intron and 1-kb areas from both ends of the expressing intron were defined as the relevant region. For “**Exon skipping**”, the 3' to 5' ends of the overlapping exons at both ends of the skipping and the surrounding 1 kb region were defined as the relevant areas, respectively. “**Cassette exons**” were counted only if they used one different exon each; other complex structures were classified as “**Complicated exon**”. The 3' to 5' ends of the overlapping exons at both ends of the cassette exon and a 1 kb area from each end were defined as the relevant region.
