## Additional_file2 for "Identification of Region-Specific Gene Isoforms in the Human Brain Using Long-Read Transcriptome Sequencing and Their Correlation with DNA Methylation"

**Command lines used in this study:**

**Iso-Seq Analysis Using pbBioConda**

#-------------------- Primer Removal & Demultiplexing

lima --isoseq --dump-clips --peek-guess -j 32 ./$sample/$sample.hifi_reads.bam ./$sample/isoseq_primers.fasta ./$sample/$sample.hifi.demult.bam

#-------------------- Trimming PolyA Trails & Concatemer Removal

isoseq3 refine --require-polya ./$sample/$sample.hifi.demult.5p--3p.bam ./$sample/isoseq_primers.fasta ./$sample/$sample.hifi.flnc.bam

#-------------------- Cluster

isoseq3 cluster ./$sample/$sample.hifi.flnc.bam ./$sample/$sample.hifi.polished.bam --verbose --use-qvs

#-------------------- Mapping

cp /[directory_1]/hg38.fasta [directory_2]/$sample/

pbmm2 align ./$sample/hg38.fasta ./$sample/$sample.hifi.polished.hq.bam ./$sample/$sample.hifi.aligned.bam -j 32 --preset ISOSEQ --sort --log-level INFO

#-------------------- Collapse

isoseq3 collapse ./$sample/$sample.hifi.aligned.bam ./$sample/$sample.hifi.collapsed.gff

**SQANTI3**

conda activate SQANTI3.env

#-------------------- QC

export PYTHONPATH=$PYTHONPATH: [directory_for_cDNA_Cupcake]/

python sqanti3_qc.py $sample.hifi.collapsed.gff gencode.v41.chr_patch_hapl_scaff.annotation.gtf /

hg38.fasta -d $sample.SQ3 -o $sample.SQ3 --CAGE_peak [directory_3] /human.refTSS_v3.1.hg38.bed /

--polyA_motif_list [directory_3]/mouse_and_human.polyA_motif.txt --polyA_peak [directory]/atlas.clusters.2.0.GRCh38.96.bed /

-n 16 --saturation --report both --isoAnnotLite --gff3 Homo_sapiens_GRCh38_RefSeq_78.gff3 -fl $sample.hifi.collapsed.abundance.txt

#-------------------- Filtering (Rule filter)

python sqanti3_filter.py rules $sample.SQ3_classification.txt -j [jsonfile: filtering.json] -o $[output_name] -d $[output_directory]

#-------------------- Filtering (Machine learning filter)_ Used only for the purpose of filtering ISM in the subsequent analysis

python sqanti3_filter.py ML $sample.SQ3_classification.txt -d $[output_directory]

**TALON**

#-------------------- Database construction

talon_initialize_database --f [directory]/gencode.v41.annotation.gtf --g hg38 --a gencode41 --o myDatabase

#-------------------- Preparation for each sample file for TALON analysis

minimap2 -ax splice -uf --secondary=no -C5 -ax splice:hq [directory]/hg38.fasta -t30 \ $sample_final.fasta > $sample_final.hg38.sam

samtools calmd $sample_final.hg38.sam [directory]/hg38.fasta/

--output-fmt sam > $sample_final.hg38.MDtagged.sam

#-------------------- Running TALON

talon --f config.csv/

--db myDatabase.db /

--build hg38 /

--cov 0.95 /

--identity 0.95 /

--o [output_file_name]

talon_create_GTF --db=myDatabase.db /

--annot=gencode41 /

-b hg38 /

--observed /

--o=[output_gtf_filename]

#-------------------- Output data creation

\# R program:

x<-read.table('[output_file_name]_read_annot.tsv',sep='/t',header=T)

x.$sample <- subset(x, dataset=="$sample")

write.table(x.$sample '$[output_filename_2].tsv', quote=F, row.names=F, sep='/t')

#-------------------- Merge with the SQANTI3 output file, the classification file

\# R program:

x<-read.table('[output_filename_2].tsv',sep='/t',header=T)

y <- read.table('$sample_classification.txt',sep='/t',header=T)

y$read_name <- y$isoform

y <- y[, colnames(y) != "X"]

y <- y[, colnames(y) != "X.1"]

m <- merge(x, y, by='read_name')

m$superID <- paste(m$annot_gene_id, m$annot_transcript_id, sep = "_")

write.table(m, '$sample.talon.classification.tsv', quote=F, row.names=F, sep='/t')
