## Additional_file3 for "Identification of Region-Specific Gene Isoforms in the Human Brain Using Long-Read Transcriptome Sequencing and Their Correlation with DNA Methylation"

**R code:**

#-------------------- Cluster analysis

x<-read.table('[data_file]', sep='/t', header=T, row.names=1)

x2 <- t(x)

d <- dist(x2)

ans <- hclust(d, method="ward.D2")

plot(ans)

#-------------------- PCA

prcomp3 <- function(df)

{

df <- na.omit(df)

ans <- prcomp(df)

ans$loadings <- t(t(ans$rotation)*ans$sdev)

ans$eigenvalues <- ans$sdev^2

print.default(ans)

invisible(ans)

}

x<-read.table('[data_file]', sep='/t', header=T, row.names=1)

x2 <- t(x)

ans <- prcomp3(x2)

pos <- c(1,1,1,1,2,2,2,2,3,3,3,3)

add_data_2 <- data.frame(pos)

names(add_data_2) <- c("pos")

df_out <- as.data.frame(ans$x)

data <- cbind(df_out, add_data_2)

plot(data$PC1,data$PC2,pch = 21, cex=1 ,bg = c(2, 3, 4)[unclass(data$pos)],lwd=0,ylim = c(-40,40),xlim=c(-40,40))

text(data$PC1,data$PC2,colnames(x),pos=3, cex=1)

#-------------------- t test

x<-read.table('[data_file]', sep='/t', header=T, row.names=1)

t.test(x$value~x$group)

#-------------------- Wilcoxon rank-sum test

x<-read.table('[data_file]', sep='/t', header=T, row.names=1)

wilcox.test(x$value~x$group)

#-------------------- Chi-square test, Fisher’s exact test and calculation of ORs (Repeat for multiple lines of data)

x <- read.table("[data_file]",header=TRUE)

nrow <- nrow(x)

sink("[output_filename]")

for (i in 1:nrow){

dat1_1 <- x[i,2:3]

dat1_2 <- x[i,4:5]

dat1 <- merge(dat1_1, dat1_2)

if ((dat1_1[1,]>0) || (dat1_2[1,]>0)) {

dat2 <- matrix(unlist(dat1), ncol=2, byrow=TRUE)

table <- as.table(dat2)

ans <- prop.test(table)

chi <- ans$statistic

pvalue <- ans$p.value

ans2 <- fisher.test(table)

pvalue_2 <- ans2$p.value

rate1 <- dat1_1[1]/(dat1_1[2]+dat1_1[1])

rate2 <- dat1_2[1]/(dat1_2[2]+dat1_2[1])

#OR

x <- table

odds <- (x[1,1]/x[1,2]) / (x[2,1]/x[2,2])

#OR (95%CI)

y1 <- log((x[1,1]/x[1,2]) / (x[2,1]/x[2,2]))

y2 <- 1/x[1,1] + 1/x[1,2] + 1/x[2,1] + 1/x[2,2]

interval <- exp(y1 + qnorm(c(0.025,0.975)) * sqrt(y2))

result <- sprintf("%.3e,%.3e,%.3f,%.3e,%.3e,%.3f,%.3f,%.3f,%.1f,%.1f,%.1f,%.1f",rate1,rate2,chi,pvalue,pvalue_2,odds,interval[1],interval[2],dat1_2[1],dat1_2[2],dat1_1[1],dat1_1[2])

print(result,quote=F,row.names=F)

}

}

sink()

#-------------------- MeDeCom

library(MeDeCom)

Data <- read.table("[methylation_datafile]")

Data2 <- as.matrix(Data)

medecom.result<-runMeDeCom(Data2, 2:10, c(0,10^(-5:-1)), NINIT=10, NFOLDS=10, ITERMAX=300, NCORES=5)

pdf("[directory]/plotParameters.pdf")

plotParameters(medecom.result)

dev.off()

pdf("[directory]/plotParameters2.pdf")

plotParameters(medecom.result, K=8, lambdaScale="log")

dev.off()

lmcs<-getLMCs(medecom.result, K=8, lambda=0.00001)

str(lmcs)

write.table(lmcs, "[directory]/lmcs.txt")

pdf("[directory]/dendrogram.pdf")

plotLMCs(medecom.result, K=8, lambda=0.00001, type="dendrogram")

dev.off()

prop<-getProportions(medecom.result, K=8, lambda=0.00001)

str(prop)

write.table(prop, "[directory]/prop.txt")

pdf("[directory]/Proportions.pdf")

plotProportions(medecom.result, K=8, lambda=0.00001, type="barplot")

dev.off()

pdf("[directory]/heatmap.pdf")

plotProportions(medecom.result, K=8, lambda=0.00001, type="heatmap")

dev.off()
